# Chemistry of cation hydration and conduction in a skeletal muscle ryanodine receptor

**DOI:** 10.1101/732172

**Authors:** Zhaolong Wu, Congcong Liu, Hua Yu, Duan Kang, Yinping Ma, Xuemei Li, Lei Zhang, Chun Fan, Xin-Zheng Li, Chen Song, Chang-Cheng Yin, Youdong Mao

## Abstract

Ryanodine receptors (RyRs) are Ca^2+^-regulated Ca^2+^ channels of 2.2-megadalton in muscles and neurons for calcium signaling. How Ca^2+^ regulates ion conduction in the RyR channels remains elusive. We determined a 2.6-Å cryo-EM structure of rabbit skeletal muscle RyR1, and used multiscale dynamics simulations to elucidate cation interactions with RyR1. We investigated 21 potential cation-binding sites that may together rationalize biphasic Ca^2+^ response of RyR1. The selectivity filter captures a cation hydration complex by hydrogen-bonding with both the inner and outer hydration shells of water molecules. Molecular dynamics simulations suggest that adjacent Ca^2+^ ions moving in concert along ion-permeation pathway are separated by at least two cation-binding sites. Our analysis reveals that RyR1 has been evolved to favor its interactions with two hydration shells of cations.

---

Calcium is an important second messenger—an intracellular signaling ion—that regulates a wide variety of cellular activities, such as neurotransmission, secretion, fertilization and cell migration (*1*). The release of Ca^2+^ ions from intracellular stores is predominantly mediated by two related Ca^2+^ channel families: the ryanodine receptors (RyRs) (*2-5*), the largest known ion channels of approximately 2.2-MDa molecular weight, and inositol 1,4,5-trisphosphate receptors (*6*). RyRs are high-conductance, monovalent- and divalent-conducting channels that are regulated by multiple factors, including Ca^2+^, Mg^2+^, ATP, phosphorylation, redox active species, and by their interactions with regulatory proteins such as FKBPs and calmodulin (*5, 7*). In most tissues, RyR channels are primarily activated by the influx of Ca^2+^ via plasma-membrane Ca^2+^ channels, resulting in a Ca^2+^-induced rapid release of Ca^2+^ from intracellular storage such as endoplasmic reticulum. In mammalian skeletal muscle, RyR1 channels are mechanically activated by direct interaction with voltage-gated L-type Ca^2+^ channel dihydropyridine receptor (DHPR or Ca_V_1.1) on the plasma membrane (*5, 8, 9*).

The general architectures of RyRs have been revealed by cryogenic electron microscopy (cryo-EM) in both basal and activated states (*10-18*). These structures provided a basis for understanding the regulatory mechanisms of RyRs activation and gating. However, many outstanding questions remain unanswered. How do cations interact with RyR channels? What is the hydration structure of cations permeating through the RyR channels? How do RyRs catalyze the translocation of cations and selectively conduct Ca^2+^ over high levels of background monovalent cations? To address these questions, we determined the cryo-EM structures of rabbit skeletal muscle RyR1 in two distinct closed states (designated state **1** and state **2**) and an open state (designated state **3**) at nominal resolutions of 2.6, 3.3 and 6.3 Å, respectively (Fig. 1, figs. S1-3, Table S1), under the same buffer condition in the presence of 10 µm Ca^2+^ and in the absence of other RyR1 regulatory proteins, such as immunophilins FKBP12/FKBP12.6 that were used in previous studies (*12, 14, 15*). The imaged RyR1 was complexed with the soluble domain of the auxiliary protein junctin, a transmembrane regulator of RyR1 that may stimulate channel opening (*19, 20*) (fig. S1). Unfortunately, in all three cryo-EM maps, we did not observe the density of the soluble domain of junctin, probably because it was bound to a flexible region near the outer surface of RyR1. Thus, the interaction between RyR1 and junctin remains unresolved. Structural comparisons among the three states and with other available RyRs structures reveal a conformational transition pathway that reproduces previous structural findings (*11-13, 17*) (figs. S4, S5).

**Figure 1.**
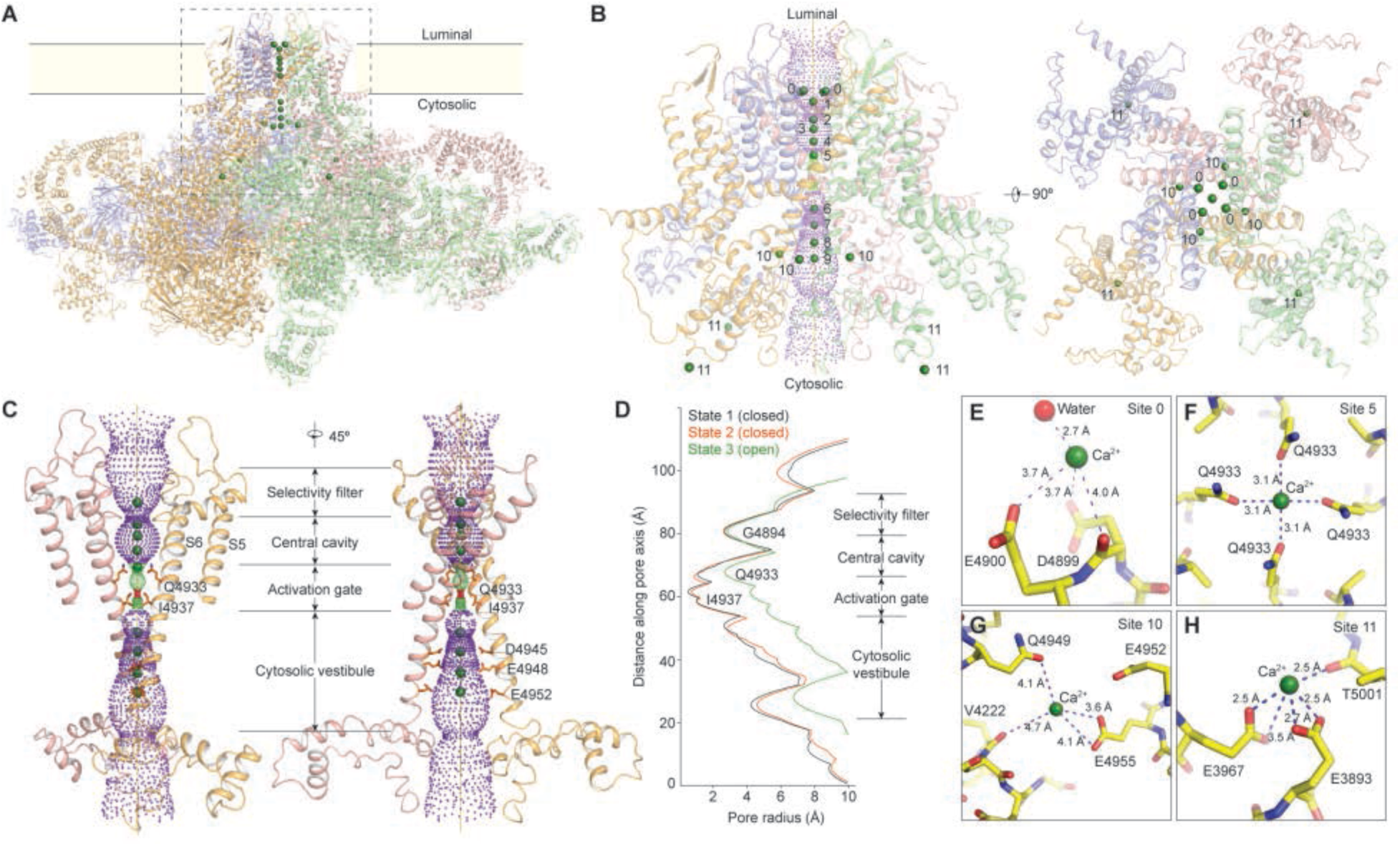
Mapping of 21 potential cation-binding sites in the RyR1 channel. (**A**) Overview of the cation-bound RyR1 tetramer structure in state **1** at 2.6-Å resolution determined by cryo-EM. (**B**) Close-up views of the 21 potential cation-binding sites in the RyR1 channel in two orthogonal perspectives. The sites are labeled with sequential numbers. The symmetric sites are designated with the same numbering. (**C**) Rendering of the central channels in two perspectives rotated 45° vertically. Channel pore domains are shown as cartoons. The solvent-accessible surface of the central channel was estimated using the program HOLE (*36*). For clarity, two opposite subunits of RyR1 are omitted. (**D**) Plots of the pore radius in three states calculated by the program HOLE (*36*). (**E**-**H**) Close-up views of Ca^2+^ interactions with oxygen atoms from side-chains and main-chains at Site 0 (**E**), Site 5 (**F**), Site 10 (**G**) and Site 11 (**H**). In all panels **A**-**C**, the protein structure is presented in transparent cartoons, and Ca^2+^ ions in green spheres.

The cryo-EM map of RyR1 tetramer in state **1** exhibits 21 local density peaks that are potentially cation-binding sites (Fig. 1A, B, Movie S1). The rationality of the cation assignment into these density peaks was extensively examined by combining *ab initio* quantum mechanical density functional theory (DFT) calculations with classical molecular dynamics (MD) simulations (figs. S6, S7, Movie S2). Nine density peaks reside along the ion permeation pathway in state **1** (Fig. 1B, C), which we name Sites 1-9 from the luminal to the cytosolic side. By contrast, we only observed three density peaks corresponding to Sites 4, 5 and 7 in the cryo-EM map of state **2** (fig. S5A). In state **1**, each density peak at Sites 1, 4 and 7 is surrounded by eight islets of density peaks forming a square antiprism cage, which were tentatively modelled as water molecules that form an inner hydration shell directly coordinating the cation at the center (Figs. 2-4) (*21-23*). Three additional density peaks compatible with bound cations were also observed per protomer (Sites 0, 10, 11; Fig. 1E-H). Among all potential cation-binding sites, the strongest ion density is located at Site 11 and embraced by Glu3893 and Glu3967 from the core solenoid domain (residues 3667-4174) and Thr5001 from the C-terminal domain (CTD, residues 4957-5037) (Fig. 1H). Site 11 is the only Ca^2+^-binding site that has been identified previously in RyR1 structures (*12*).

**Figure 2.**
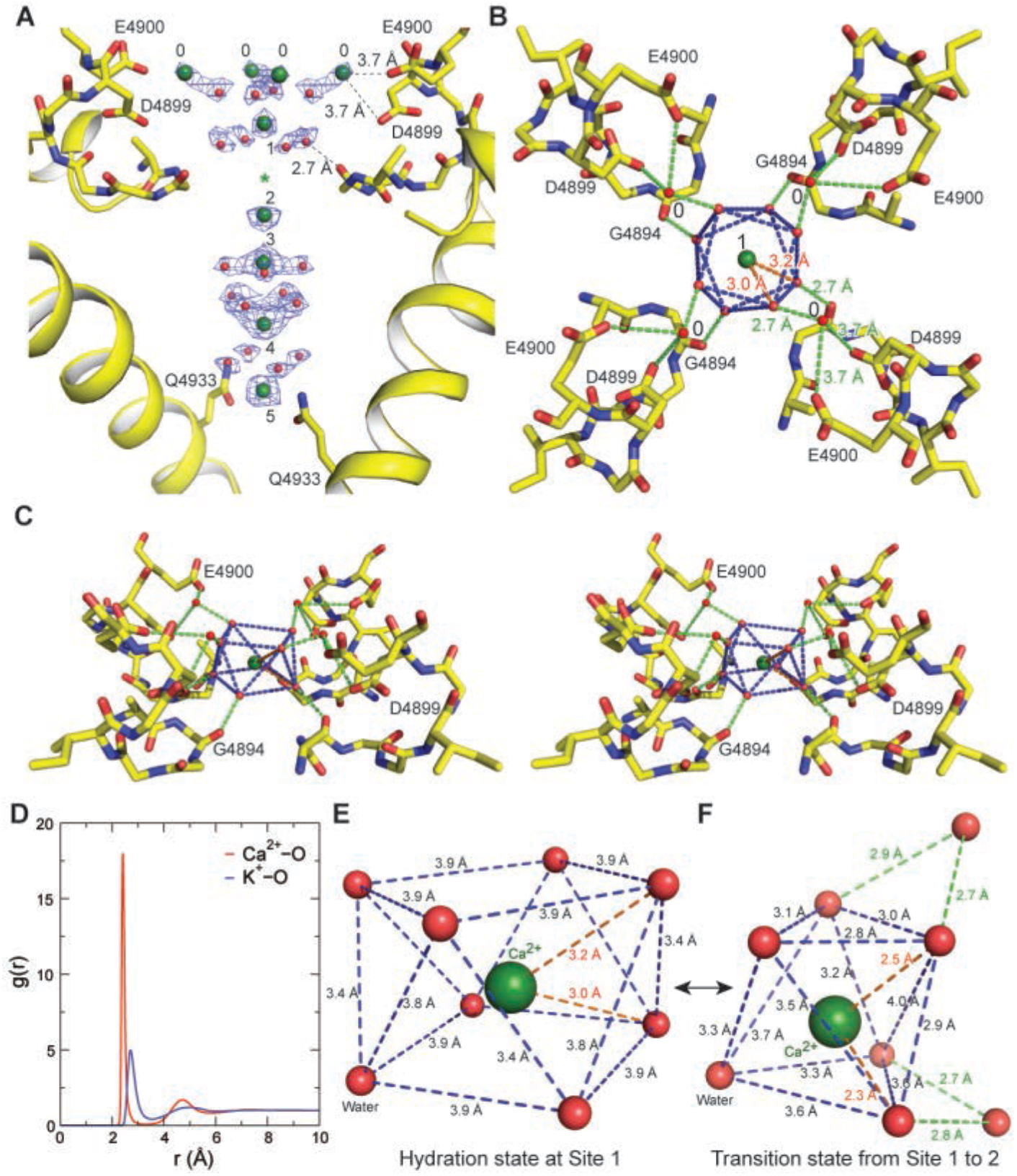
Recognition of cation hydration complex by the selectivity filter. (**A**) Close-up lateral view of cation-binding Sites 0-5 from the selectivity filter to the central cavity in state **1**. The cryo-EM densities of cations and water molecules directly coordinating the cations are shown as blue meshes at 2σ level. For clarity, two opposite subunits of RyR1 are omitted. (**B**) Top view of the cation hydration complex bound at Site 1, with Site 0 occupied by water molecules mediating hydrogen bonds with negatively charged residues. (**C**) Stereoscopic view of the cation hydration complex at Site 1. (**D**) The radial distribution functions (RDFs) of water oxygen atoms around cations from our MD simulations. r, the distance between water oxygen and cation. The first (left) and second (right) peaks correspond to the inner and outer hydration shells of water molecules, respectively. The outer hydration shell of K^+^ is significantly weaker than that of Ca^2+^. (**E**) Detailed structure of the Ca^2+^ hydration complex with only the inner hydration shell shown, determined by cryo-EM. (**F**) Detailed structure of the Ca^2+^ hydration complex in the transition state when passing the narrowest point (marked by a green asterisk in panel **A**) of the selectivity filter, determined by *ab initio* DFT calculations. Ca^2+^ ions and oxygens atoms are shown as green and red spheres, respectively. Blue and green dashed lines mark the atom distances among the inner and outer shell of waters of Ca^2+^ hydration, respectively.

RyR1 features a unique selectivity filter that comprises a very short passage of constriction, in sharp contrast to other calcium channels of known structures (*22, 24-27*) (Fig. 2A). The lower four water molecules in the inner hydration shell of the cation at Site 1 appear to form hydrogen bonds with the upward-pointing carbonyl oxygen atoms in Gly4894 at the selectivity filter (Fig. 2B, C). It is energetically unfavorable for cations to occupy both Sites 0 and 1 simultaneously due to electrostatic repulsion. When a cation occupies Site 1, water molecules can occupy Site 0 and constitute part of the outer hydration shell of the cation, which mediates a hydrogen-bond network with carboxyl oxygen atoms in Asp4899 and Glu4900 (Fig. 2B, Movie S3). Our MD simulations indicate that a single cation hydration complex can carry two hydration shells of water molecules that diffuse together with the cation, although the outer hydration shell is less stable than the inner hydration shell (Fig. 2D). It is also noteworthy that the outer hydration shell of K^+^ is much less stable than that of Ca^2+^, so is the inner hydration shell (Fig. 2D). It appears that the selectivity filter indirectly interacts with a cation by forming hydrogen bonds with both the inner and outer shells of water molecules in the cation-hydration complex (Fig. 2A, B). We reason that such a mode of cation recognition may favor multivalent cations over monovalent ones, because a cation with less stable hydration shells, such as K^+^, could suffer from lower affinity and specificity (fig. S7F, G). Further, our *ab initio* DFT simulations show that single Ca^2+^ hydration complex permeates through the constriction of the selectivity filter by moving two water molecules from the inner to outer hydration shell and with the inner hydration shell transiently rearranging into an octahedron of six water molecules (Fig. 2E, F, fig. S6A), whereas K^+^ can keep five water molecules in the inner hydration shell when crossing over the selectivity filter at Gly4894 (fig. S6B). Taken together, our data may help retionalize how RyR1 selects Ca^2+^ over K^+^ or Na^+^ by a factor of ∼7-8 folds (*28, 29*).

Inside the central cavity, four density peaks (Sites 2-5) compatible with cations are observed along the channel axis in state **1** (Figs. 1B, C, 3A). To understand the functional role of these cation-binding sites with respect to the ionic occupancy state during permeation, we performed targeted MD simulations in which the activation gate was steered from state **1** (closed) to state **3** (open) (fig. S7C). The results suggest that only one Ca^2+^ ion can enter the central cavity at a time in state **1**, and it prefers to stay at Site 4, consistent with the observation of the strongest on-axis cryo-EM density peak at this site in state **1** (Fig. 2A, fig. S5A). Eight water molecules on the vertical position of Site 3 appear to be stabilized into an octagon constituting the second hydration shell of Ca^2+^ in the central cavity. Impressively, the locations of these stabilized water molecules from our MD simulations agree with their assignment in the cryo-EM density (fig. S7B). In state **3**, with channel opening, Site 5 becomes the preferred cation-binding site before the ion passes the gate (Fig. 3C). Therefore, the cation-binding sites in the transmembrane pore are state-dependent. Nonetheless, in all MD-simulated events, two nearest neighboring Ca^2+^ ions moving in concert along the ion permeation pathway—from the selectivity filter to the activation gate—are separated by at least two cation-binding sites or approximately 11 Å (Fig. 3C, D), in line with the observation that the two strongest on-axis cryo-EM density peaks before the gate are at Sites 1 and 4 in state **1**, or at Sites 1 and 5 in state **3** (fig. S5A). These results together support an ion-permeation mechanism that the Ca^2+^ hydration complexes knock off one another without breaking their innermost hydration shells of water molecules.

**Figure 3.**
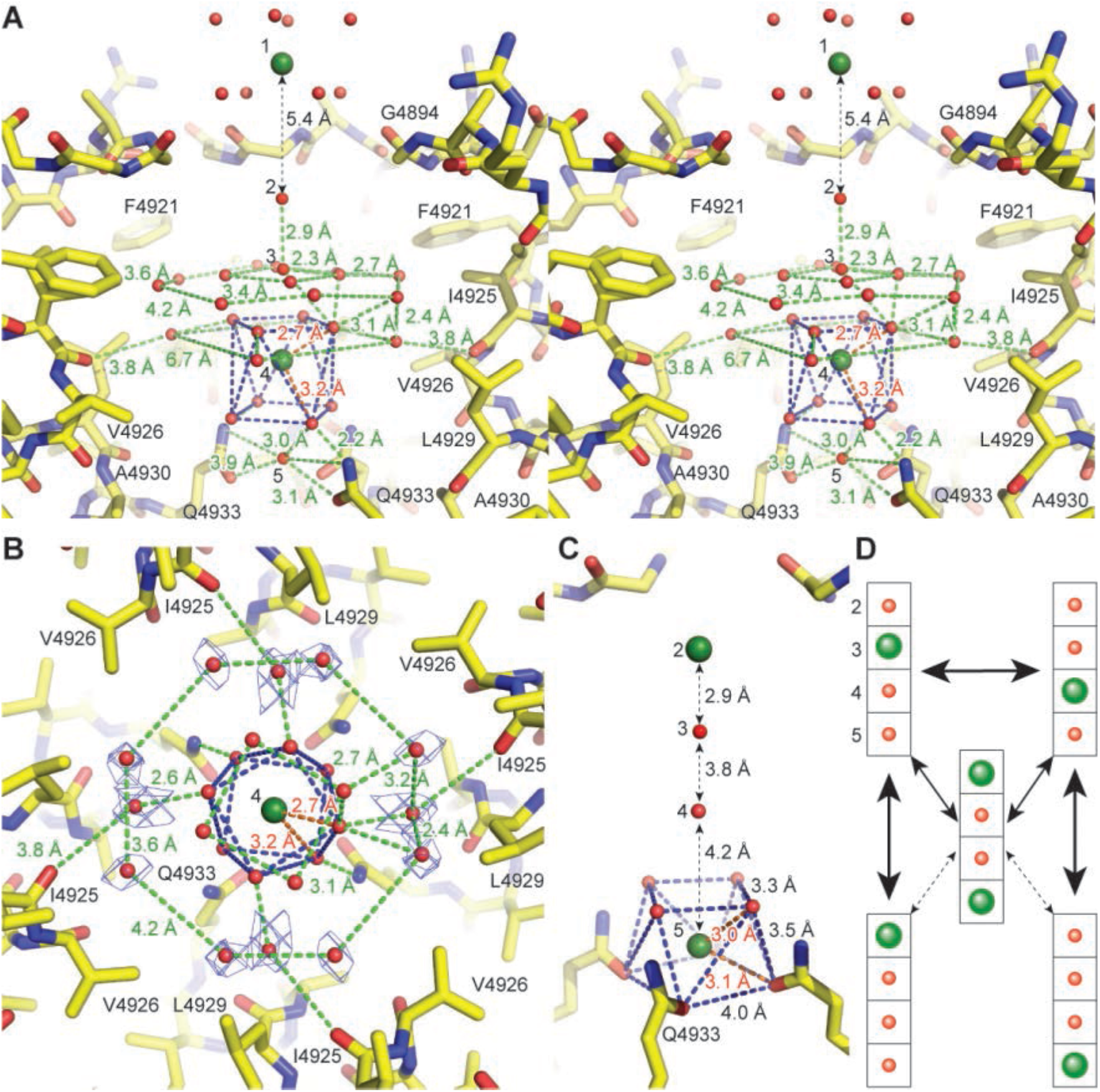
Interactions of cation hydration complex with the central cavity and mechanism of ion permeation. (**A**) Stereoscopic view of the cation hydration complex inside the central cavity. For clarity, one subunit of RyR1 is omitted. The inner hydration shell of cation at Site 4 is highlighted with blue dashed lines between water molecules. 20 water molecules were tentatively modelled into the surrounding density peaks inside the transmembrane cavity, forming a hemispherical hydrogen-bond network. When Site 1 is occupied by Ca^2+^, water molecules instead occupy Sites 2 and 3, manifesting a plausible occupancy state as suggested by our MD simulations. Blue and green dashed lines mark the atom distances among the first and second shell of waters of Ca^2+^ hydration, respectively. (**B**) Top view of the cation hydration complex at Site 4 inside the central cavity. The cryo-EM densities of water molecules not directly coordinating Ca^2+^ are shown as blue meshes at 2σ level. (**C**) Possible alternative occupancy state where Sites 2 and 5 are taken by Ca^2+^ ions, and Sites 3 and 4 by water molecules. (**D**) An ionic occupancy state diagram of RyR1 in the central transmembrane pore, derived from cryo-EM data and MD simulations, shows the proposed cation permeation mechanism.

The constriction at the activation gate of RyR1 is narrowed by Gln4933 and Ile4937 in states **1**-**3**. Both K^+^ and Ca^2+^ cations are energetically forbidden for permeation across the gate in state **1** (fig. S6C, D). By contrast, our *ab initio* DFT and classical MD simulations suggest that a Ca^2+^ ion can carry eight water molecules when it travels through the Ile4937-narrowed constriction in state **3** (fig. S6E). However, a K^+^ ion may lose two water molecules during permeation through this constriction (fig. S6F) (*21, 23*). These differences imply that the activation gate may also contribute to the Ca^2+^ selectivity by imposing dehydration energy penalty on K^+^ ions. In support of this finding, previous single-channel experiments have demonstrated that mutation of Asp4938 to asparagine results in a significant reduction in Ca^2+^ selectivity over K^+^ ions (*29, 30*). In state **1** structure, four equally spaced density peaks from Site 6 to 9 are lined along the channel axis in the cytosolic vestibule (Fig. 4A, fig. S5A). The average distance of adjacent sites is 4.9 ± 0.1 Å. The outer hydration shell of the cation at Site 7 connects the Ca^2+^ ion through a spherical hydrogen-bond network to the side-chain nitrogen or oxygen atoms in three charged residues Arg4944, Asp4945 and Glu4948 from four pore-lining S6 helices (Fig. 4B, C, Movie S5). Our *ab initio* DFT calculations suggest that Ca^2+^ binding at Site 7 may stabilize the closed state of RyR1 by holding the four S6 helices together below the activation gate (Fig. 4A-C, fig. S6G, H). Notably, the hydrogen-bond network structure entrapping a cation at Site 7 in state **1** is not compatible with state **3** conformation, where Asp4945 and Glu4948 are translated ∼4 Å away from the channel axis (Fig. 4D, E).

**Figure 4.**
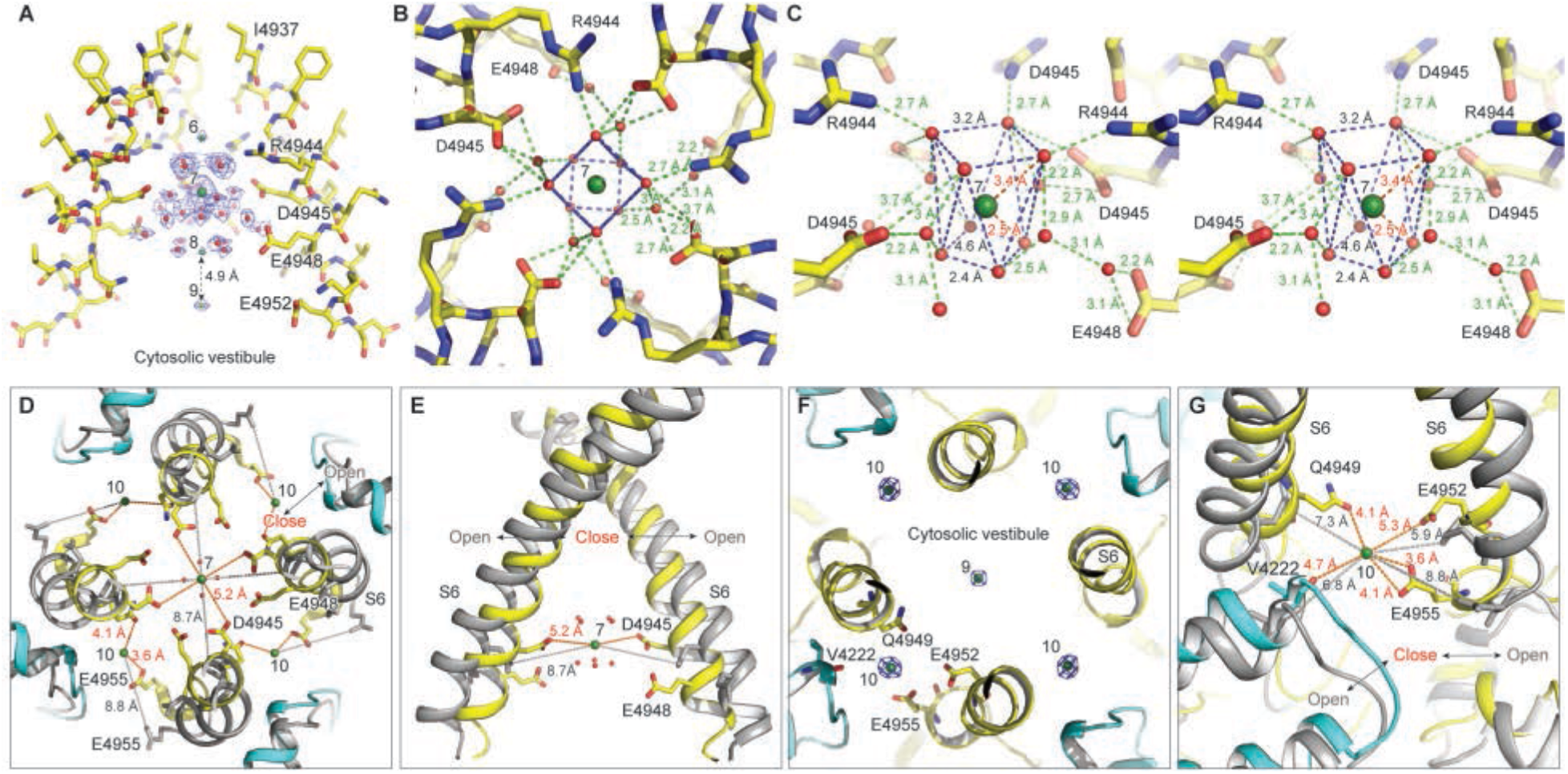
Interactions of cation hydration complex with the cytosolic vestibule. (**A**) Close-up side view of potential cation-binding sites 6-9 in the cytosolic vestibule in state **1**. The cryo-EM densities of cations and water molecules coordinating the ions are shown as blue meshes at 1.8σ level. (**B**) Top view of the cation hydration complex at Site 7 inside the cytosolic vestibule. The inner hydration shell of Ca^2+^ ion at Site 7 is highlighted with blue dashed line between water molecules coordinating Ca^2+^. Other hydrogen bonds outside of the inner hydration shell are shown as green dashed lines. (**C**) Stereoscopic view of the cation hydration complex at Site 7. Blue and green dashed lines mark the atom distances among the inner and outer shells of water molecules of cation hydration, respectively. For clarity, one subunit is omitted in **A** and **C**. (**D)**, Top-view structural comparison of Sites 7 and 10 inside the cytosolic vestibule between the closed (color) and open (grey) states from a perspective along the channel axis. (**E**) Side-view structural comparison of Site 7 inside the cytosolic vestibule between the closed (color) and open (grey) states. For clarity, two opposite subunits of RyR1 are omitted. (**F**) Top view of the cryo-EM densities at Sites 9 and 10 with narrowed depth, shown as blue meshes at 1.8σ level. The Sites 9 and 10 are located at the approximately same horizontal position along the channel axis. (**G**) Structural comparison around Site 10 at the entry of the cytosolic vestibule between the closed (color) and open (grey) states.

At the entrance of cytosolic vestibule, four potential cation-binding sites with low occupancy (Site 10) in state **1** bridge the inter-subunit gaps between adjacent S6 helices and the back of the thumb and forefingers (TaF) domain directly supporting the EF-hand pair domain (Fig. 4F). Each cation at Site 10 can be coordinated with oxygen atoms in Glu4955 side-chain from one subunit, and in Gln4949 side-chain and Val4222 main-chain from the adjacent subunit (Fig. 1G). By contrast, this inter-subunit gap is ∼8 Å wider in state **3**, making it impractical for Ca^2+^ binding at Site 10 when the channel is open (Fig. 4D, G). We thus speculate that these putatively low-affinity cation-binding sites in the cytosolic vestibule (Sites 6-10) could contribute to locking the S6 helices and the TaF domain supporting EF-hand pair in the closed state by millimolar cytosolic Ca^2+^, which is compatible with previous studies (*31*). Further investigations are required to determine if these and other unknown low-affinity cation-binding sites are required for Ca^2+^-dependent inhibition of RyRs.

To understand how Ca^2+^ dynamically regulate the RyR1 activity, we performed stochastic dynamics (SD) simulations of Ca^2+^ release via RyR1. The gating dynamics of RyR1 are described by the Fokker-Plank equation as a nonequilibrium, stochastic diffusion process on the experimental free-energy landscape (fig. S8A). The RyR1 dynamics are allosterically regulated by the cation-RyR1 interactions that are composed of both repulsion and attraction forces (*32*). The repulsion force can be exerted by hydrophobic interactions between Ile4837 and Ca^2+^ hydration complexes permeating through the gate. The attraction force can be mediated by the electrostatic interactions of Ca^2+^ hydration complexes at Sites 6-10 in the cytosolic vestibule (Fig. 4). Given the tight activation gate in state **3**, the free energy of permeating cation hydration complex may be transferred in part to the channel by removal of water molecules from the outer hydration shell or by inelastic collision of the hydration complex with the gate. This is expected to result in an increase in the probability of channel opening (fig. S8A). This hypothesis is in line with recent experiments showing the enhancement of RyR1 activation by multivalent cations in the permeation pathway (*33*), and is further supported by the quantitative agreement of our SD simulations with single-channel experiments on the biphasic cytosolic Ca^2+^ response curves (fig. S8, Table S2) (*34, 35*). To sum up, our structural and computational studies have unraveled unexpected mechanisms of Ca^2+^ selectivity and permeation, Ca^2+^-regulated gating of RyR1. The observation of extensively rich cation-binding sites in RyR1 reveals a unique set of design principles of calcium channel architecture, and suggests that RyRs may have been evolved to favor their interactions with two hydration shells of cations.

## Acknowledgments

The authors thank S. Zhang, Y. Zhu, D. Yu, J. Xu for assistance in data collection, NVIDIA corporation for accessing their proprietary DGX-2 system in data processing. This work was funded in part by National Natural Science Foundation of China grant 91530321 (Y.M.), 31570732 and 31770785 (C.C.Y.), 21873006 (C.S.), Natural Science Foundation of Beijing Municipality grant Z180016 (Y.M.), National Key Research and Development Programs 2017YFA0504702 (C.C.Y.), 2016YFA0500401 (C.S.), 2016YFA0300900 and 2017YFA0205003 (X.Z.L.) in Ministry of Science & Technology of China. The cryo-EM experiments and data collection were performed at Electron Microscopy Laboratory and Cryo-EM Platform at Peking University. The data processing was performed in part in the Weiming No. 1 and Life Science No. 1 High-Performance Computing Platform at Peking University. Molecular dynamics simulation was performed in part on the High-Performance Computing Platform of the Center for Life Sciences at Peking University.

## Author contributions

Y.M., C.C.Y., C.S. and X.Z.L. devised this study. C.L. and L.Z. purified the protein complexes, conducted single-channel recording experiments and prepared the cryo-specimen. Z.W. and C.L. screened the condition for optimal data collection. Z.W., C.L, and X.L. collected the cryo-EM data. Z.W., C.L., Y-P.M., and C.F. processed the cryo-EM data. Z.W. and Y.M. refined the maps and the atomic models, and performed stochastic dynamics simulations. H.Y. and C.S. performed classical molecular dynamics simulations. D.K. and X.Z.L. performed ab initio quantum mechanical simulations. Y.M. integrated analyses of results from different aspects of this study and wrote the manuscript with inputs from all authors. All authors discussed the results and contributed to the manuscript preparation.

## Data availability

Cryo-EM maps and the atomic coordinates will be deposited to the Electron Microscopy Data Bank and RCSB Protein Data Bank prior to the formal publication of this manuscript.

## Material and Methods

### Purification of RyR1

RyR1 was purified from New Zealand white rabbit skeletal muscle. First, sarcoplasmic reticulum (SR) vesicles were purified (*3*). Briefly, muscle tissue (∼200 g) was minced, suspended in 500 ml of 20 mM Hepes, pH 7.4, 1 mM EDTA-Na, 0.3 M sucrose, 2 mM phenylmethylsulphonyl fluoride (PMSF), 1:1000 diluted protease inhibitor cocktail (PIC) (Sigma-Aldrich). The suspension was homogenized in a blender (3×30 s) with low and high speed respectively and centrifuged for 25 min at 8500 g, 4 °C. Then, the supernatant was centrifuged for 1 h at 110000 g, 4 °C. Afterwards, the sediment was added to 400 ml buffer containing 20 mM Hepes, pH 7.4, 0.65 M KCl, 1 mM EDTA-Na, 0.3 M sucrose, 2 mM PMSF, PIC). The solution was wobbled for 45 min on the ice. After ultracentrifugation for 35 min at 4 °C, the sediment was resuspended in 15 ml of 20 mM Hepes buffer (pH 7.4, 0.3 M sucrose, 2 mM PMSF, PIC), obtaining the SR vesicles containing ∼25 mg/ml proteins.

Next, RyR1 was purified from SR vesicles (*13*). SR vesicles was resuspended in 1 M NaCl, 20 mM Hepes, pH 7.4, 2 mM DTT, 2 mM PMSF, PIC and 1.2% CHAPS (AMRESCO)/0.6% soybean lecithin (Sigma-Aldrich). The solution was wobbled on the ice for 1 h, and then, centrifuged for 45 min at 110000 g. The supernatant was applied onto a hydroxyapatite ceramic Bio-Rad column (HA-column-I), pre-equilibration by 200 mM NaCl, 10 mM Hepes, pH 7.4, 0.5% CHAPS/0.25% soybean lecithin, 2 mM DTT, 2 mM PMSF, PIC. After prewashing, proteins were eluted with 15 ml, 200 mM NaCl, 10 mM Hepes, pH 7.4, 0.5% CHAPS/0.25% soybean lecithin, 2 mM DTT, 2 mM PMSF, protease inhibitors cocktail, 200 mM K_2_HPO_4_. The collected eluate was loaded on a 100-kDa cut-off Amicon centrifugal filter (Millipore) and centrifugation at 1000 g, 4 °C. The obtained protein solution was then applied on a 5%∼20(w/v) linear sucrose gradient and centrifugation for 16 h in a Beckman SW28 rotor at 26000 rpm. Afterwards, proteins were applied on the HA-column-II. For the size-exclusion chromatography (20 mM Hepes, 200 mM NaCl, 0.5% CHAPS, 2 mM DTT, 2 mM PMSF, PIC), Superose 6 Increase column was used and the RyR1-enriched solution was collected, concentrated, rapidly frozen in liquid nitrogen, and stored at - 80°C.

### Purification of junction

Recombinant junctin (protein id=AAF37204.1) from rabbit was expressed and purified by Ni-NTA His Bind Resin. Briefly, Escherichia coli Bl21 (DE3) cells containing the codon-optimized gene of junctin, with the transmembrane region deleted, were grown at 37 °C to OD_600_ of 0.4∼0.6. Junctin was induced for expression by adding 1mM IPTG at 22 °C for 16 h. Cells were harvested by centrifugation at 4500 g at 4 °C for 30 min. The sediment was re-suspended by 50 mM Na_2_HPO_4_, pH 7.4, 150 mM NaCl and lysed by sonication. The mixture was centrifuged to remove the cell debris. The protein solution was applied on the Ni-NTA column. After prewashing, junctin was eluted by the lysis buffer containing 50 mM imidazole. The pooled fractions were concentrated and applied on a size-exclusion chromatographic column (SUPERDEX^TM^ 75 10/300 GL). The junctin-enriched fractions were pooled and rapidly frozen by liquid nitrogen to be used in the subsequent experiments.

### Surface plasmon resonance

To examine the binding of purified junctin with the purified RyR1, RyR1 complexes were connected to the biotin, acting as the stationary phase. Different concentrations of purified junctin (0 nM, 0.8 nM, 1.6 nM, 3.2 nM, 6.4 nM, 12.5 nM, 25 nM, 50 nM, 100 nM, 200 nM) acted as the mobile phase. Time for association and dissociation are 180 s and 600 s, respectively. The experiment was carried out in Biacore^TM^ T200.

### Density gradient centrifugation for the interaction of RyR1 and junction

RyR1 and junctin reacted in the buffer containing 100 mM NaCl, 20 mM Hepes, pH 7.4, 10 μM Ca^2+^, 2 mM DTT, 2 mM PMSF, PIC and 0.5% CHAPS/0.25% soybean lecithin. The molar ratio of RyR1 and junctin is 1:8. After wobbled on the ice for 1 h, the reaction mixture were loaded on the top of the 5%∼20(w/v) linear sucrose gradient and centrifugation for 16 h in a Beckman SW28 rotor at 26000 rpm. Fractionated into 1.5 ml fractions, the resulting samples were tested by SDS-PAGE. The fractions containing both RyR1 and junctin were pooled together to load on a PD-10 column (GE) in buffer composed of 100 mM NaCl, 20mM Hepes, pH 7.4, 10 μM Ca^2+^, 2 mM DTT, 2 mM PMSF, PIC to remove CHAPS and soybean lecithin. The collected solution of RyR1-junctin complex was concentrated to about 0.4 mg/ml for preparing cryo-EM sample.

### Single-channel recordings

Purified RyR1 was reconstituted into microsomes as described(*37*). Briefly, the purified RyR1 was brought to 5 mg/ml lipid mixture (5:3 PC:PE) and placed in a dialysis bag (14 kDa MWCO) and dialyzed against 1 liter of dialysis buffer (0.5 M NaCl, 0.1 mM EGTA, 0.2 mM Ca^2+^, 1 mM DTT, 2 mM PMSF and 20 mM Hepes-Na, pH 7.4) to remove free detergent. Afterwards, samples were sub-packaged, snap-freezing in liquid nitrogen and stored. Microsomes were fused to planar lipid bilayers formed by painting a lipid mixture of phosphatidylethanolamine and phosphatidylcholine (Avanti Polar Lipids) in a 3:1 ratio in decane across a 150-μm hole in cups (Warner Instruments) separating 2 chambers (*12*). Symmetric solutions (250 mM KCl, 20 mM Hepes, pH 7.4) were used to record channel currents. A −80 mV potential bias was applied (from *trans* to *cis* chamber) during single-channel recording experiments. The concentration of free Ca^2+^ in the *cis* chamber was calculated with Windows Maxc program (v2.5.1, http://web.stanford.edu/~cpatton /downloads.html). Electrical signals were filtered at 2 kHz through an eight-pole low-pass Bessel filter (Warner Instruments), and digitized at 4 kHz. Data acquisition was performed using Digidata 1440A, Axopatch 200B and Clampex 10.3 software (Molecular Devices). The recordings were analyzed by using Clampfit 10.3 and Graphpad software.

### Sample preparation for the cryo-EM

Aliquots of 5 µl of RyR1-junctin complex (∼0.4 mg/ml) were placed on glow-discharged R2.0/2.0 holey carbon grids (GIG), on which a homemade continuous carbon film had previously been deposited. Grids were blotted for 2 s and flash-frozen in liquid ethane using an FEI Vitrobot. Cryo-grids were transferred to an FEI Titan Krios electron microscope that was operating at 300 kV and equipped with the post-column Gatan BioQuantum energy filter connected to Gatan K2 Summit direct electron detector. Data were collected semi-automatically using SerialEM (*38*) at a magnification of 105,000 times in super-resolution electron counting mode on a K2 Summit direct electron camera with the Gatan BioQuantum operating in the zero-loss imaging mode (20-µm energy slit). The calibrated physical pixel size is 1.37 Å, and the super-resolution pixel size is 0.685 Å on the specimen scale. A movie of 40 frames was recorded in 10 s per exposure, with a total dose of 49 e/Å^2^. A total of 10,009 movies were collected from the Titan Krios electron microscope within two sessions of data collection.

### Cryo-EM data processing and reconstruction

The raw frames of 10,009 movies were gain-corrected for their gain reference and were drift-corrected to generate EM micrographs by MotionCor2 program (*39*). Each micrograph was used for the determination of the defocus parameters according to its Thon rings with Gctf program (*40*). After auto-picked and verified using RELION 2.1 (*41*) and EMAN2 software (*42*), 835,709 particles of RyR1 were extracted from 9,528 good micrographs for the following analysis. All reference-free 2D and 3D classifications were carried out in RELION 2.1/3.0 (*41*) and ROME 1.1 (*43*), which combined the regularized maximum-likelihood based image alignment and manifold learning-based classification. 3D refinement that refined the Euler angles and x/y-shifts of each particle to further improve the resolution of density maps was completed in RELION. Both 3D classification and refinement of maps were done at a pixel size of 1.37 Å, which was binned by two folds from the raw data in the super-resolution mode.

After several rounds of iterative 2D and 3D classifications, bad particles and non-particle contaminants were removed from the dataset of RyR1 particles, and 529,554 high-quality particles were left. First, all good particles were aligned to one consensus model to determine their angular and shift parameters. The initial model was low-pass filtered to 60 Å to eliminate template bias in the reference of 3D calculation. Then focused 3D classification was performed to determine the heterogeneity of particles. A mask including the transmembrane region of the consensus model was created using program RELION, which was applied to do 3D classification by skipping image alignment. This process made it possible to detect subtle conformational changes of the transmembrane domain. By setting class number K = 10 in the focused 3D classification, which both show the closed channel core domain but differ largely in the location of peripheral cytosolic domains. To test if our dataset containing particles with open-channel state, a multi-reference 3D classification with 3D alignment skipped was performed, using three input reference maps from low-pass filtered the maps of state **1**, state **2** and the previously published open state (EMD-8378) (*12*) to 5 Å. By this approach of supervised 3D classification without 3D alignment, 3,922 particles of the open state were obtained. Finally, high-resolution refinement of each map was done by using the same input reference of our consensus model low-pass filtered to 60 Å, with a global mask applied and C4 symmetry imposed, resulting in three distinct conformational states, named state **1**, state **2** and state **3**.

In order to improve the local resolution of density maps, the CTF parameters of each individual particle were refined locally with program Gctf (*40*). Based on the x/y-shift and orientation parameters of each particle from the high-resolution 3D refinement, we reconstructed two half-maps of each state using raw single-particle images at super-resolution mode with a pixel size of 0.685 Å by ROME software (*43*). The global resolutions of the finial reconstructions of states **1**, **2** and **3** were 2.6, 3.3 and 6.3 Å respectively, measured by the gold-standard Fourier shell correlation (FSC) at 0.143-cufoff on two independently refined half-maps. For the sake of visualization, an estimated negative B-factor was applied to all structures to sharpen their density maps, and local resolution variations were then calculated by rome_res module in the ROME software (*43*) on two half-maps of each state that implemented and parallelized the same algorithm in ResMap (*44*).

### Atomic model building and refinement

To build the initial atomic model of RyR1 in state **1**, we use previously published structure (PDB ID 5T15) as a starting model and then manually improved the main-chain and side-chain fitting in Coot (*45*). To fit the model to the density map, we first conducted rigid fitting of the segments of the model in Chimera (*46*), and then rebuild the mismatched main-chain and side-chain manually in Coot. The ligands, including Ca^2+^ ions and water molecules, were fitted manually in Coot, and verified by either quantum chemistry or MD simulations (see below). Atomic model refinement of the fitted models was done in real space with Phenix (*47*), with NCS constraint. The refined atomic model of state **1** was then used as starting reference for real-space refinement of state **2** in Phenix (*47*). Although the channel core domain of state **3** is virtually identical to the Ca^2+^/ATP/Caffeine-bound open-state RyR1 structure, our state-**3** map lacks the densities for the auxiliary transmembrane helices (TMx) and for the immunophilins FKBP12/FKBP12.6 (also called calstabin1/calstabin2), consistent with the density maps of both states **1** and **2**. We thus revised the published atomic model (PDB ID 5TAL) of the Ca^2+^/ATP/Caffeine-bound open-state RyR1 structure by deleting segments or chains without corresponding densities in our state-**3** map and use it as the starting reference for real-space refinement in Phenix (*47*) with both NCS and secondary structure constraints.

### Quantum mechanical simulations

The quantum mechanical density functional theory (DFT) calculations were performed with the Vienna *ab initio* Simulation Package (*48*) (VASP), using projector augmented wave (PAW) pseudopotentials (*49*) and an energy cutoff of 700 eV for the expansion of the wave functions. The Perdew–Burke–Ernzerhof (PBE) functional was employed to address the exchange-correlation interacctions (*50*). To account for the long-range dispersion interactions, the DFT-D2 approach by Grimme was adopted (*51*). Our focus of these DFT calculations is to see how the static energies of the system and the structures of the hydrated ions change upon pushing the ions through the channel. In the optimizations the thresholds for the electronic convergence criterion and ionic relaxation were set to 1 × 10^-5^ eV and 5 × 10^-3^ eV, respectively. Considering the computation cost, local structures of the activation gate in state **1** and state **3**, the selectivity filter and the potential well at Site 7 in state **1** were employed in the simulations, and their structures were given in fig. S6. The Ca^2+^/K^+^ ions were hydrated with 4 H_2_O in the state-**1** activation gate simulation, and 8 H_2_O in other simulations. The simulated systems were computed with periodic images separated by at least 10-Å vacuum layer, and the periodic box sizes of 25 × 25 × 25 Å^3^, 30 × 30 × 25 Å^3^, 35 × 35 × 25 Å^3^, and 30 × 30 × 28 Å^3^ were used for the activation gate in state **1** and state **3**, the selectivity filter and the potential well at Site 7 in state **1**, respectively. The *k*-space integration of the simulation box was done with a single Γ point. The atoms near the pathway of Ca^2+^/K^+^ and H_2_O were relaxed in the calculations.

### Molecular dynamics simulations

GROMACS 2018.6 was used to conduct the coarse-grained and all-atom molecular dynamics simulations (*52*). The targeted molecular dynamics (TMD)(*53*) simulations were performed with the PLUMED plugin (*54*). The coarse-grained self-assembly molecular dynamics simulation with the Martini2.2 force field (*55*) and elastic network (*56*) was performed to construct the simulation systems. Initially, water and POPC (1-palmitoyl-2-oleoyl-sn-glycero-3-phosphocholine) molecules were randomly placed in the simulation box around the coarse-grained pore domain (residue ID: 4820-4956) of the state-**1** structure. A 200-ns production simulation was conducted to obtain a coarse-grained system under NPT ensemble (isothermal-isobaric ensemble), after energy minimization (200,000 steps) and a preliminary NVT (canonical ensemble) equilibration (50 ps) with the position restraint applied on the heavy atoms of the protein with a force constant of 10^3^ kJ/mol/nm^2^. The periodic boundary condition was used. The time step was 20 fs. The velocity-rescale algorithm (*57*) with a time constant of 1.0 ps and the Berendsen algorithm (*58*) with a time constant of 10 ps were utilized to maintain the temperature at 320 K and the pressure at 1.0 Bar, respectively. The reaction-field method with a cut-off of 1.1 nm was used to calculate electrostatics, and the same cut-off was used to calculate the van der Waals interactions. The last frame of the coarse-grained trajectory was extracted, in which the transmembrane domain of RyR was well-embedded in a POPC bilayer, and then converted (*59*) into an all-atom system within the CHARMM36 force field (*60, 61*). Then, two all-atom systems were constructed respectively, by aligning and embedding (*62*) the pore domain of the state-**1** structure with and without the cryo-EM-determined Ca^2+^ and water molecules into the bilayer, and 0.15 M CaCl_2_ was added to neutralize the system. The all-atom molecular dynamics simulations were conducted to obtain the stable binding sites of the Ca^2+^ in the pore domain of the cryo-EM structure based on an improved calcium model (*63*). The CHARMM36m force field (*64*) with TIP3P water model (*65*) was used. A 500-ps NVT equilibration and a 500-ps NPT equilibration were performed after energy minimization. Then, two 500-ns production simulations were conducted and the trajectories of the last 300 ns were used for data analysis. Position restraints with a force constant of 1000 kJ/mol/nm^2^ were applied on the backbone atoms of the protein for both the equilibration and the production simulations. The periodic boundary conditions were used and the time step was 2 fs. The velocity-rescale algorithm (*57*) with a time constant of 0.5 ps was used to maintain the temperature at 310 K. Protein, membrane, water and ions were coupled separately. The Parrinello-Rahman algorithm (*66*) with a time constant of 5 ps was used to maintain the pressure at 1.0 Bar. The Particle-Mesh Ewald (PME) method (*67*) was used to calculate electrostatics and the van der Waals interactions were considered within a cut-off of 1.2 nm.

We used PLUMED 2.6 (*54*) in combination with GROMACS 2018.6 to perform the TMD simulations to model the opening process of the pore from state **1** to state **3**. The starting system of the TMD simulation was obtained from the last frame (500 ns) of the all-atom molecular dynamics simulation performed above. The pore domain of the open-state structure (state **3**) of RyR1 was used as the target structure. Three independent 200-ns TMD simulations in the presence and absence of a transmembrane potential of 200 mV were performed, respectively. For all the TMD simulations, the harmonic restraints were applied to control the RMSD (root mean square deviation) of the protein backbones between the simulated trajectory and the target structure. The Kearsley algorithm (*68*) (TYPE = OPTIMAL) was used to calculate the alignment matrix for further calculation of the RMSD between the target and the instantaneous configuration of the steered structure. The force constant of the harmonic restraint was increased linearly from 0 to 10^6^ kJ/mol/nm^2^ in the first 1 ns and then maintained at this value for the rest of the simulations. The other simulation parameters were the same as in the conventional all-atom molecular dynamics simulations described above. The trajectory and density analysis was performed with VMD (*69*).

### Reconstitution of energy landscape

In order to analyze the dynamical behavior of the RyR1 channel, we first need to know its free energy landscape, which can be obtained from the cryo-EM data (*70*). After 2D classification, all remaining particles were randomly separated into 11 subsets, and subsequent 3D classification in every subset reconstructed 53 density maps in total. After density maps reconstructed by particles fewer than 1300 were removed and the channel region of RyR1 was masked, 33 cryo-EM reconstructions were left for further dimensionality reduction by either principal component analysis (PCA) or t-distributed stochastic neighbor embedding (t-SNE). When we reduced the dimensions of 3D map data by PCA to 2, we found all points almost distributed along a straight line (fig. S8D). Thus we straightforwardly reduced the dimension to one, and mapped the first reaction coordinate to the channel radius 𝑅 at the constriction of activation gate in our high-resolution structures to calculate the free energy function 𝐺 = 𝐺(𝑅), based on the Boltzmann distribution. For any pair of cryo-EM reconstructions 𝑖 and 𝑗, the Boltzmann distribution suggests that the difference of free energy ΔΔ𝐺 between them is

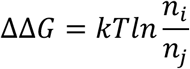

where 𝑘 is the Boltzmann constant, 𝑇 is the absolute temperature, and 𝑛_*i*_, 𝑛_*j*_. is the number of particles used to reconstruct the cryo-EM density maps 𝑖 and 𝑗, respectively. For simplicity, the free energy of the reconstruction 𝑖 can be valued as

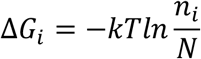

where 𝑁 is the total particle. A constant that can be added to the free energy for any cryo-EM reconstruction is omitted here. As shown in fig. S8A, the free-energy function has two local minima that correspond to the closed and open states, respectively. Consistent with this results, the t-SNE-derived pseudo-energy landscape shows a circular valley corresponding to state **1**, which is separated by a circular barrier to shallow wells corresponding to state **3** (fig. S8E). Note that the one-dimensional free-energy profile plotted against the radius of the gate constriction ignores the conformational changes in the peripheral cytosolic shell domains, whereas the two-dimensional free-energy landscape plotted against two t-SNE dimensions does not. Thus, the locations of states 1 and 2 is very close in the one-dimensional free-energy landscape (fig. S8D), but they are well separated on the two-dimensional free-energy landscape (fig. S8E).

### Stochastic dynamics simulations

To understand the gating dynamics of the transmembrane pore of RyR1, we analyzed stochastic motion of the channel interacting with hydrated calcium ions, based on the high-resolution RyR1 structure and the parameters provided by classical MD and quantum mechanical DFT simulations. As the energy well of the open state is so shallow, the free-energy profile for channel activation can be approximately simplified to a harmonic function of the gate radius *R*, which can be written as

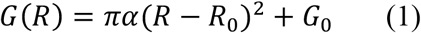

where 𝑅_0_ is the gate radius of the ground state, 𝐺_0_ is the ground energy, and 𝛼 is the area tension of channel whose order of magnitude can be derived from the experimental measurement of the free-energy landscape (Fig. 6a). Based on the theoretical model introduced previously (*32*), the Fokker-Plank equation describing the channel dynamics of RyR1 reads

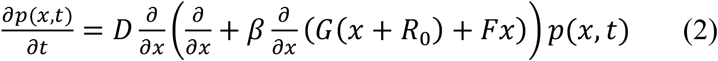

where 𝑥 = 𝑥(𝑡) = 𝑅 − 𝑅_0_ is the reaction coordinate, 𝑝(𝑥, 𝑡) is probability density at (𝑥, 𝑥 + 𝑑𝑥), 𝐷 is the diffusion coefficient, 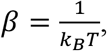 and 𝐹 is the external force exerted on the channel of RyR1. Inserting Smoluchowski approximation 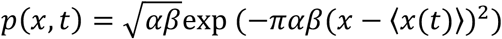 and the harmonic potential in equation (1), it is easy to derive this ordinary differential equation

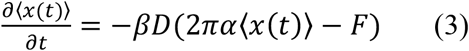

where 〈𝑥(𝑡)〉 indicates the ensemble average of the reaction coordination 𝑥(𝑡). The exact solution of equation (3) with the initial condition of 〈𝑥(0)〉 = 0 is

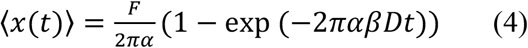

Equation (4) shows that the radius of channel changes exponentially with time 𝑡, which is consistent with the rapid-changing ions current recorded in the single-channel experiment. Let 𝑡 → ∞, when the force 𝐹 is a constant, the 〈𝑥〉 at an equilibrated state can be derived

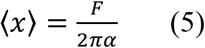

which is consistent with the result by directing inserting 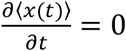 into equation (3). Equation (5) suggests that the external force 𝐹 exerted on the channel plays an important role in the change of the gate radius 𝑅. However, the force is too complicated to calculate in a precise way. In our model, two components governed by different mechanisms may contribute to the force 𝐹. The first part of force is attributed to the hydrated Ca^2+^ ions permeating through the channel of RyR1. The cryo-EM maps demonstrate that Ca^2+^ ions will bind to the negatively charged residues of the cytosolic vestibule in the closed state, whereas in the open state the density of Ca^2+^ ions in this position was not observed (fig. S5A, B). So the electrostatic interaction exerted by Ca^2+^ ions in this position stabilizes the closed state of the channel. Meanwhile, the Ca^2+^ hydration complexes exert hydrophobic repulsion to the channel when going through the constriction at the activation gate narrowed by the hydrophobic residues Ile4937. Considering the Ca^2+^ ions in the channel, this part of force is formulated as

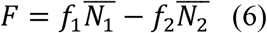

where 𝑓_1_ is the mean hydrophobic repulsion from single Ca^2+^ hydration complex outward along the radius, 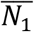 is the average Ca^2+^ ion number acting the repulsion force 𝑓_1_, 𝑓_2_ is the mean electrostatic interaction from Ca^2+^ hydration complexes inward along the radius, and 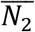 is the average Ca^2+^ ion number acting the interaction 𝑓_2_. The order of magnitude of force 𝑓_1_ and 𝑓_2_ was calculated by quantum mechanical DFT simulation (see above). Considering the probability Ca^2+^ ions bound to the protein, previous studies have suggested the relationships (*32*)

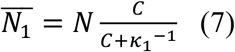

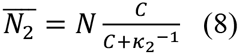

where 𝐶 is cytosolic Ca^2+^ concentration, 𝜅_1_, 𝜅_2_ is the association constants of two binding reactions, respectively, and 𝑁 is the local Ca^2+^ ion number in the channel. Inserting the function (7), (8) into (6), the mean force *F* can be written as

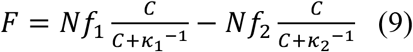

The second part of the force is regulated by allosteric activation of RyR1. The allosteric activation is structurally complicated and could change the free energy landscape in a Ca^2+^-dependent manner. In RyRs, the remote regulation via Ca^2+^ binding to sites 11 or other unknown activation sites in the cytosolic shell domains can be simplified by a modulatory coefficient function imposed on the force expressed in equation (9). So the effective mean force can be revised as

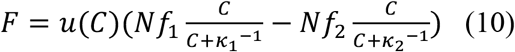

where 𝑢(𝐶) is the modulatory coefficient function of cytosolic Ca^2+^ concentrations that defines the overall sensitivity of the primed RyR1 to the Ca^2+^-dependent activation. When the cytosolic Ca^2+^ concentration is lower than certain threshold, 𝑢(𝐶) = 0 means that the channel will not be moved by any force from Ca^2+^ interactions. 𝑢(𝐶) can also assume certain value greater than 1 as mutation of some residues can allosterically stimulate the force sensitivity of the channel. As the R615C mutation (*31*) might make RyR1 to hold calcium ions more frequently in the channel, the mean force 𝐹 will increase due to the growing local ion number 𝑁 and other structural factors. Given these effects, the modulation coefficient 𝑢(𝐶) is assigned 1 and 1.42 in the case of wild-type RyR1 and R615C mutant, respectively.

In order to predict the open probability of the channel of RyR1, a radius threshold 𝑅^∗^ and 𝑥^∗^ = 𝑅^∗^ − 𝑅_0_ that calcium can be released should be defined in the model. When 𝑡 → ∞, the probability density function 𝑝(𝑥, 𝑡) can be rewritten as 𝑝(𝑥). So the open probability 𝑃_d_ can be calculated as:

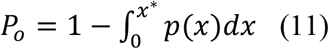

where the threshold 𝑥^∗^ was calculated by the targeted MD simulation (see above), and the probability density function 𝑝(𝑥) can be expressed as 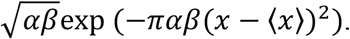 Based on our stochastic dynamics model, the mechanism that calcium ions regulate the conformational change of the channel of RyR1 was examined computationally using parameters derived by classical MD and quantum mechanical DFT calculation. To summarize, our SD modeling offers a quantitative dissection on the effects of calcium-RyR1 interactions—based on our nonequilibrium statistical mechanical model at single-channel and single-ion level, in contrast to the macroscopic model empirically hypothesizing Michaelis-Menden-type binding(*35*)— that predicts the open probability during the RyR1 activation by micromolar Ca^2+^ and inactivation by millimolar Ca^2+^ (fig. S8C, H).

---

**Fig. S1.**
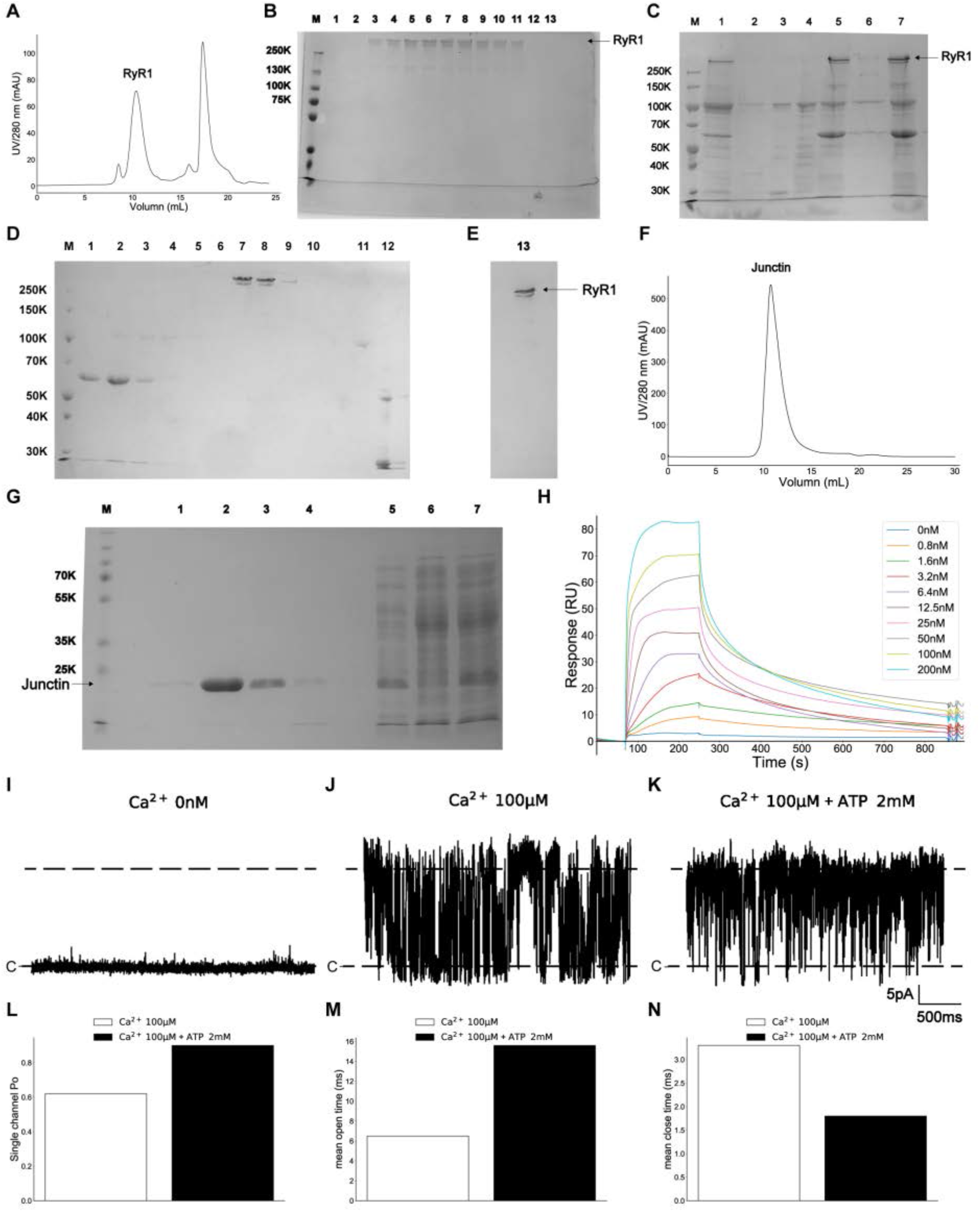
Purification and single-channel recording of RyR1 tetramer. **A**, Size-exclusion chromatography of RyR1 on Superose 6 Increase column. **B**, SDS-PAGE analysis for the RyR1 fractions collected from the size-exclusion chromatography. **C**, SDS-PAGE analysis for HA-column-I. **D**, Result of sucrose density gradient centrifugation. **E**, Final SDS-PAGE analysis of the rabbit RyR1 purified by HA-column-II. **F**, Size-exclusion chromatography of junctin. **G**, SDS-PAGE analysis for the junctin purified by gel-filtration chromatography. **H**, Results of surface plasmon resonance (SPR) to verify the binding of purified junctin with the purified RyR1. (**I**-**K**) Results of RyR1 single-channel recording. The *cis* chamber contains 0 mM free Ca^2+^ (**I**), 100 µM free Ca^2+^ (**J**) and 100 µM free Ca^2+^, 2 mM ATP (**K**), respectively. (**L**-**N**) Single-channel open probability Po (**L**), mean open time (**M**), mean close time (**N**) with 100 µM free Ca^2+^ and 100 µM free Ca^2+^, 2 mM ATP.

**Fig. S2.**
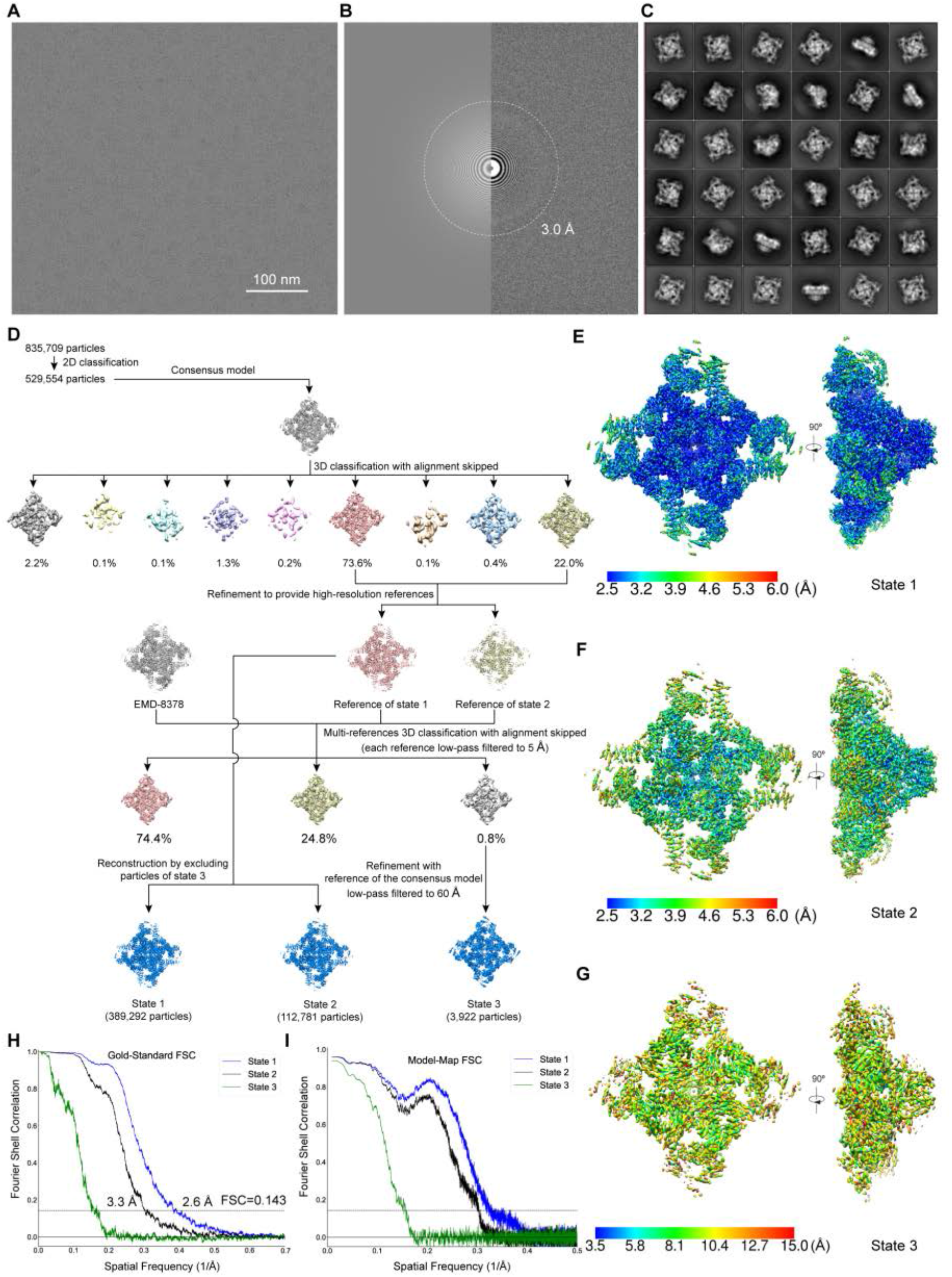
Cryo-EM structure determination of the Ca^2+^-bound RyR1 tetramer. (**A**) Typical cryo-EM micrograph of RyR1-junctin complex after drift correction. (**B**) Power spectrum evaluation of the micrograph shown in **A**. (**C**) Representative two-dimensional class averages of RyR1 particles. (**D**) Schematic showing the procedure for 3D classification and refinement of the heterogeneous dataset into three distinct conformational states for high-resolution reconstruction. An empty class of the first 3D classification is omitted in the diagram. (**E**-**G**) Local resolution estimation of states **1** (**E**), **2** (**F**) and **3** (**G**), calculated by rome_res module in ROME^50^ between two half-maps separately refined in a gold-standard procedure. (**H**) Fourier shell correlation (FSC) plots of the three maps, each calculated with two half-maps refined separately in a gold-standard procedure. (**I**) FSC plots calculated by Phenix^54^ after real-space refinement between each refined cryo-EM map and its corresponding atomic model. The color code for each state is the same in **H** and **I**.

**Fig. S3.**
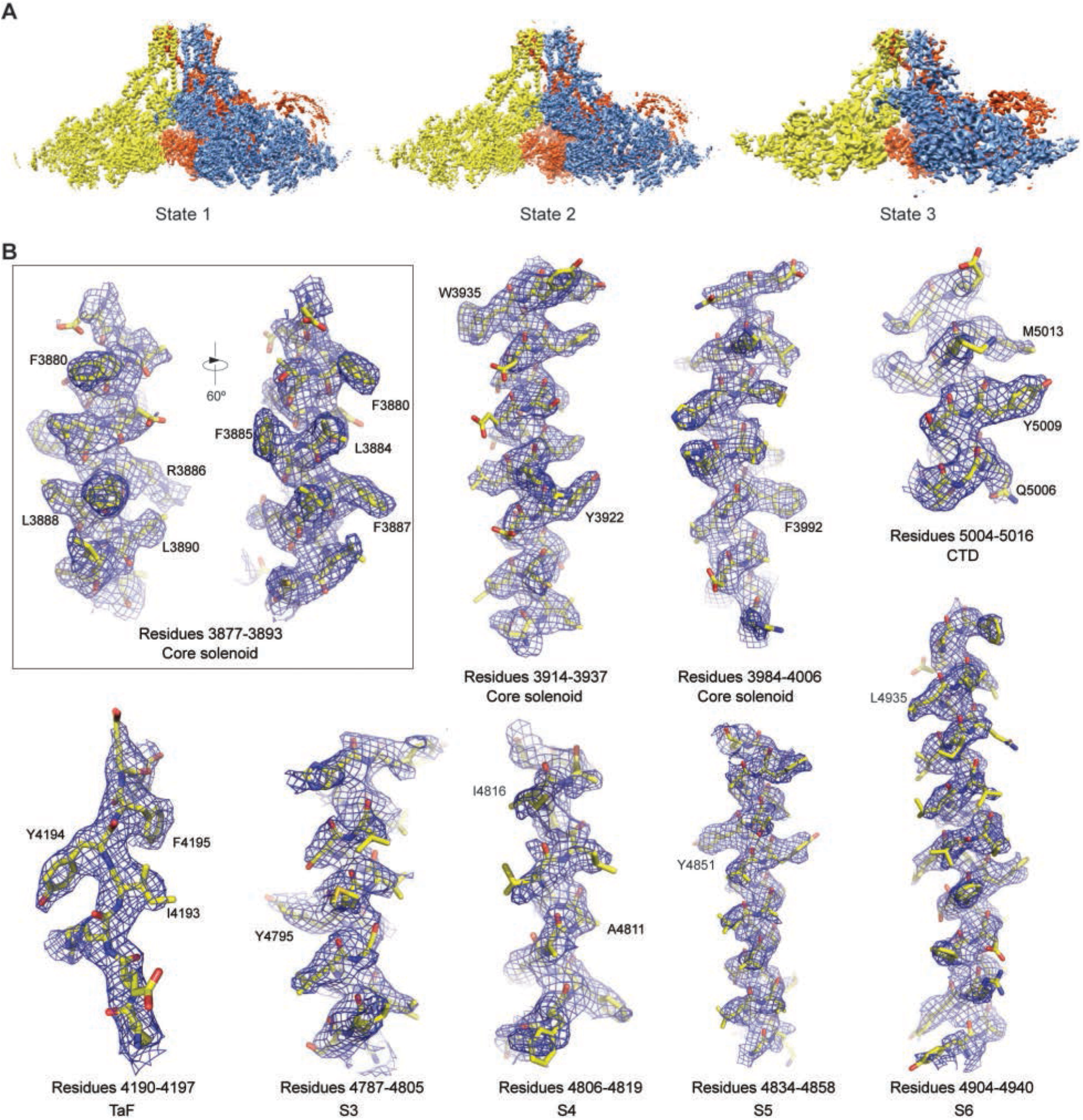
Cryo-EM density maps of RyR1 in distinct states. (**A**) Cryo-EM density maps of RyR1 in three states. One subunit is omitted to show the interior structures. (**B**) Close-up views of local high-resolution densities of secondary structures in state **1**, shown as blue meshes superimposed with the atomic models in stick representations.

**Fig. S4.**
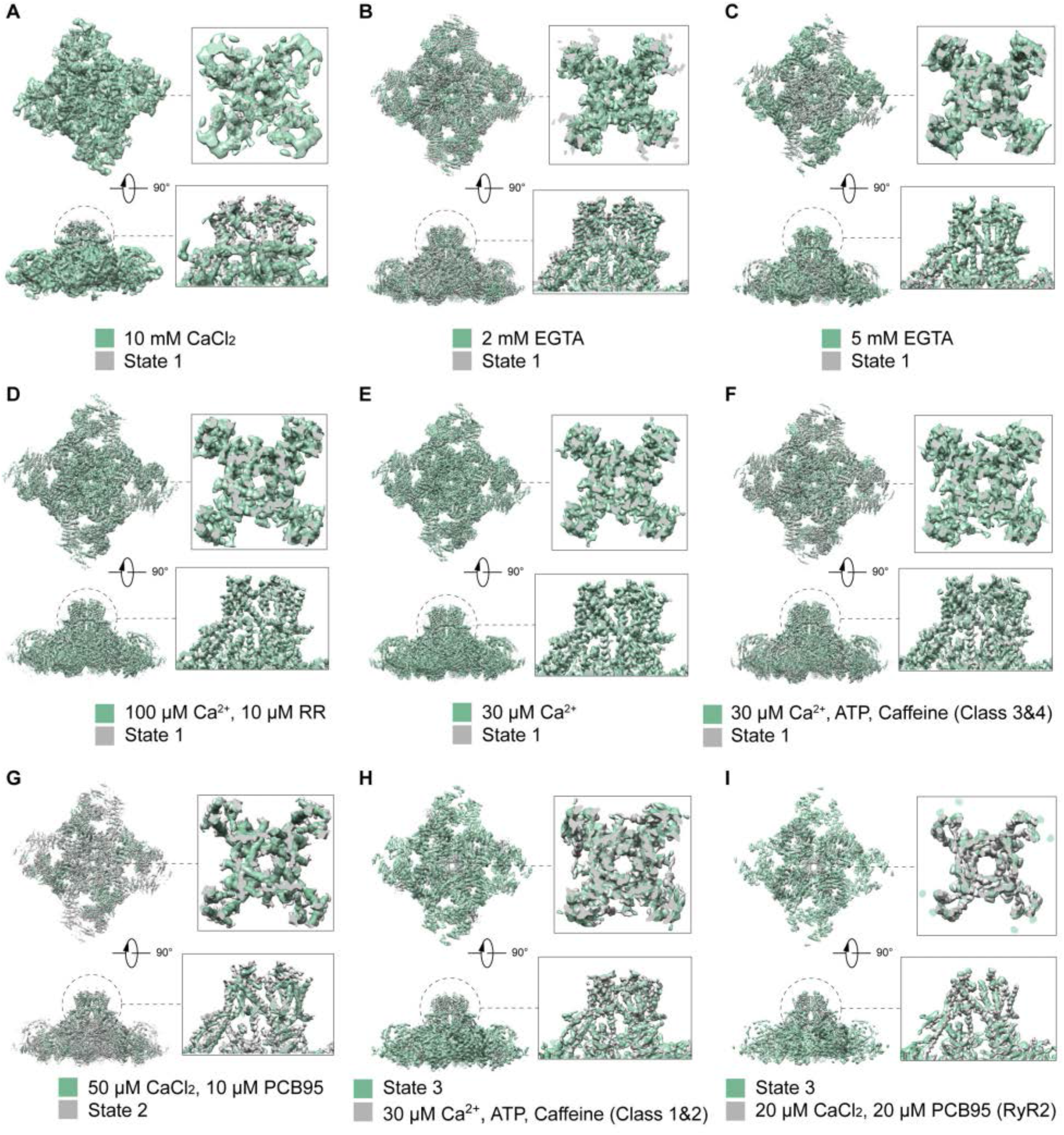
Comparisons of our RyR1 structures in different states of this study with those released cryo-EM maps of previous studies. (**A**) The state-**1** map superimposed with the EMD-2752 map (10 mM CaCl_2_)^18^. (**B**) The state-**1** map superimposed with the EMD-2807 map (2 mM EGTA)^16^. (**C**) The state-**1** map superimposed with the EMD-6106 map (5 mM EGTA)^17^. (**D**) The state-**1** map superimposed with the EMD-8073 map (100 µM Ca^2+^, 10 µM RR)^15^. (**E**) The state-**1** map superimposed with the EMD-8342 map (30 µM Ca^2+^)^14^. (**F**) The state-**1** map superimposed with the EMD-8382 map (30 µM Ca^2+^, ATP, Caffeine, Class 3&4)^14^. (**G**) The state-**2** map superimposed with the EMD-9521 map (50 µM CaCl_2_, 10 µM PCB95)^13^. (**H**) The state-**3** map superimposed with the EMD-8378 map (30 µM Ca^2+^, ATP, Caffeine, Class 1&2)^14^. (**I**) The state-**3** map superimposed with the EMD-9529 map of RyR2 (20 µM CaCl_2_, 20 µM PCB95)^19^. The 2.6-Å structure of state **1** is consistent with the cryo-EM structure of Ca^2+^-bound RyR1-FKBP12.6 complex in the ‘primed state’^14^. The channel domain structure of state **2** is very close to that of state **1**, but their peripheral cytosolic shell domains exhibit progressive dilation compared to state **1**. The open conformation of state **3**, particularly in the channel domain, is virtually identical to the cryo-EM structure of the open-state RyR1 bound with three ligands— Ca^2+^, ATP and caffeine^14^.

**Fig. S5.**
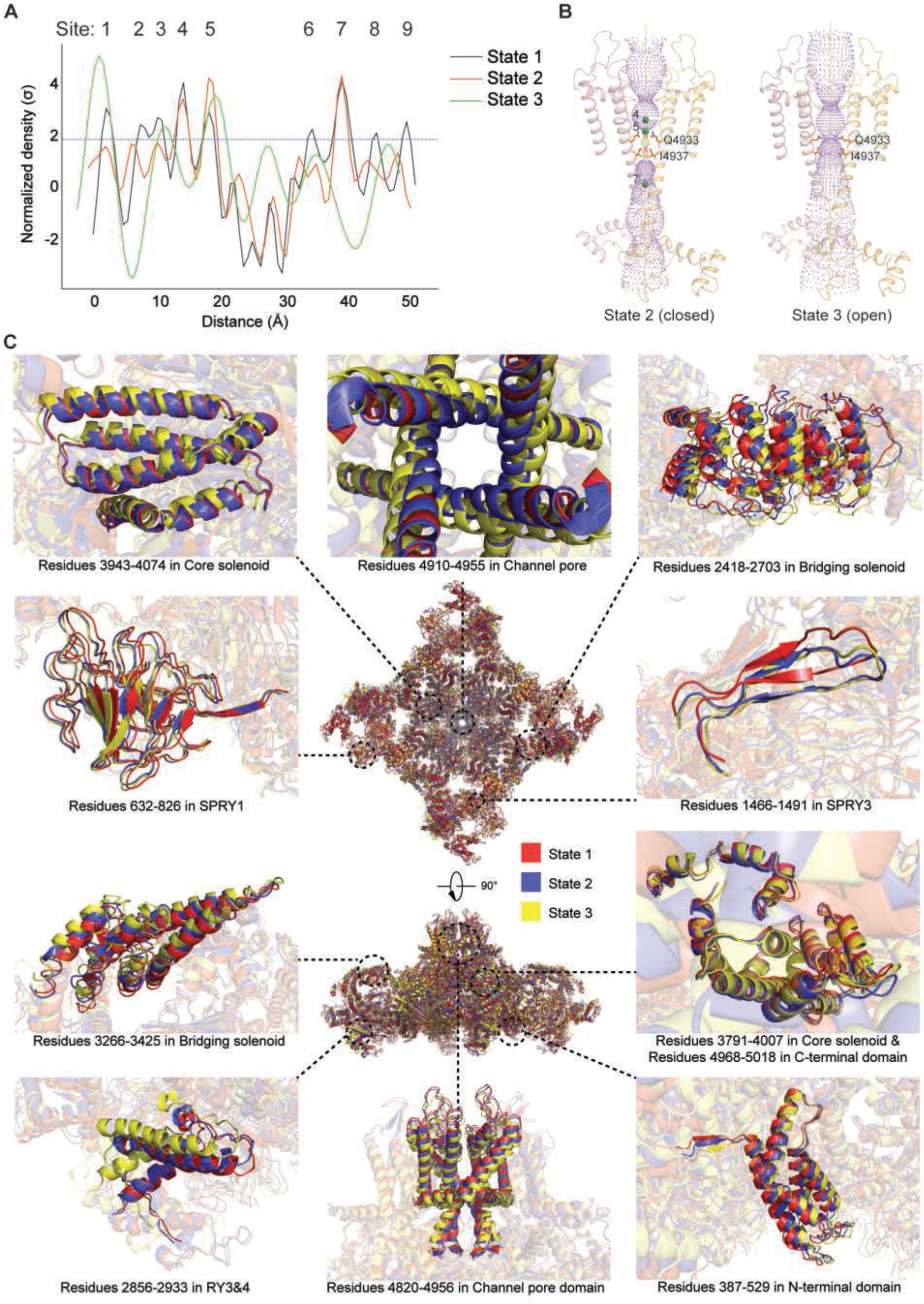
Comparison of the RyR1 structures among states 1-3. (**A**) Cryo-EM densities sampled along the channel axis show the comparison of the density peaks reflecting the ion distribution in the channel in the three states. For a fair comparison, the densities were normalized with the σ value of each map. After aligning with each other, the density values along the channel axis show consistent peaks corresponding to Ca^2+^-binding sites 1-9. The dash line marks 1.8σ. (**B**) 3D rendering of the pore radius in states **2** and **3**. The solvent accessible surface of the central channel was calculated using the program HOLE^77^. (**C**) Local structural comparison among the three states shows that the conformational changes starts from the domains in the peripheral cytosolic shell and propagates to the central channel.

**Fig. S6.**
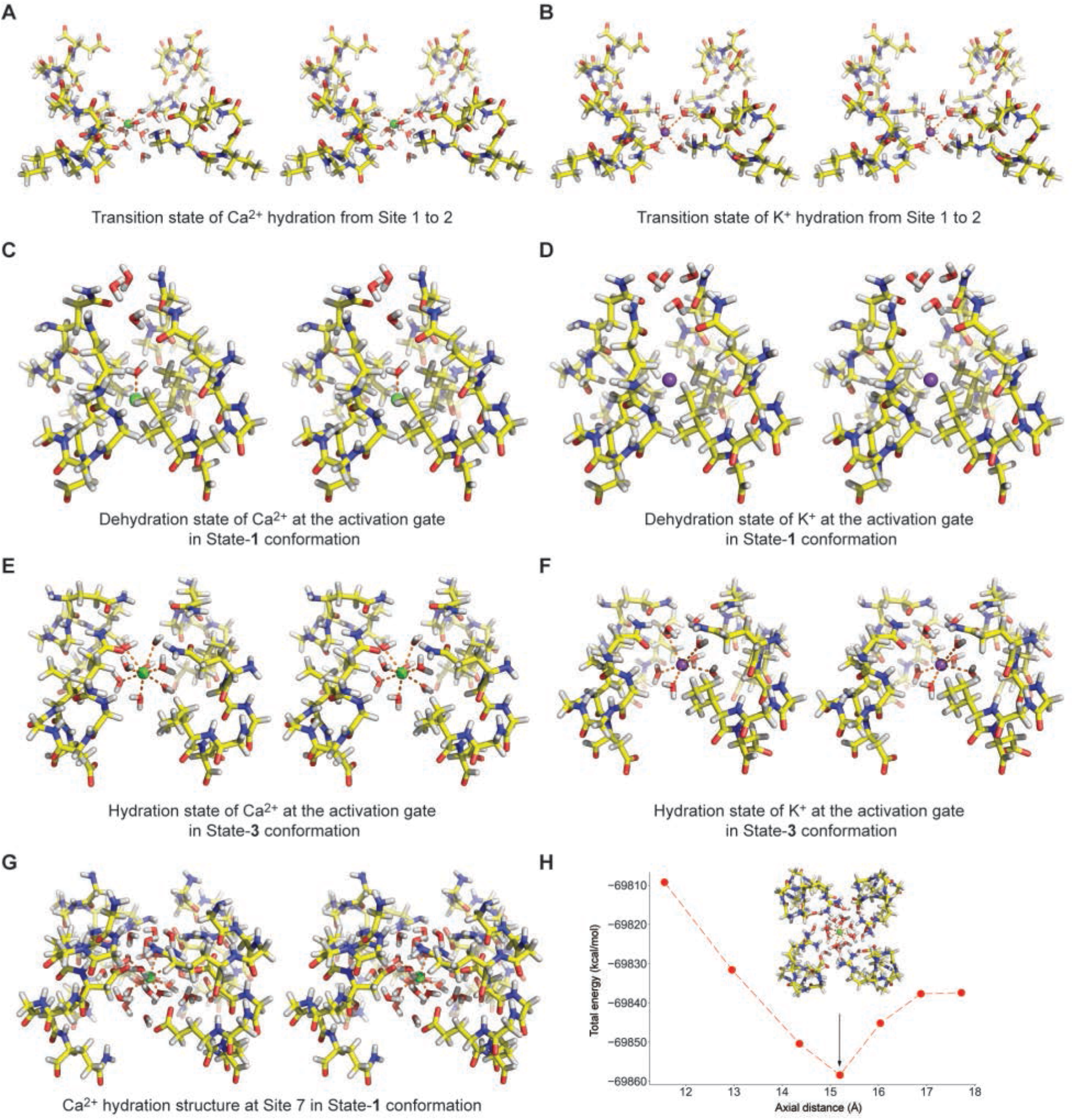
*Ab initio* DFT simulations of cation hydration and dehydration at different positions in the RyR1 channel. (**A**) Stereoscopic view of the transition state of Ca^2+^ hydration complex translating from Site 1 to Site 2 during ion conduction through the selectivity filter. (**B**) Stereoscopic view of the transition state of K^+^ hydration complex translating from Site 1 to Site 2. (**C**) Stereoscopic view of the DFT-calculated state of Ca^2+^ dehydration at the activation gate in state-**1** conformation. (**D**) Stereoscopic view of the DFT-calculated transition state of K^+^ dehydration at the activation gate in state-**1** conformation. (**E**) Stereoscopic view of the DFT-calculated state of Ca^2+^ dehydration at the activation gate in state-**3** conformation. (**F**) Stereoscopic view of the DFT-calculated transition state of K^+^ dehydration at the activation gate in state-**3** conformation. (**G**) Stereoscopic view of the DFT-calculated structure of Ca^2+^ hydration complex bound at Site 7 inside the cytosolic vestibule in state-**1** conformation corresponding to the local energy minima. (**H**) The DFT-calculated total energy of the Ca^2+^ hydration complex around Site 7 inside the cytosolic vestibule in state-**1** conformation plotted against the position along the channel axis. In all panels, Ca^2+^ and K^+^ ions are represented as green and purple spheres.

**Fig. S7.**
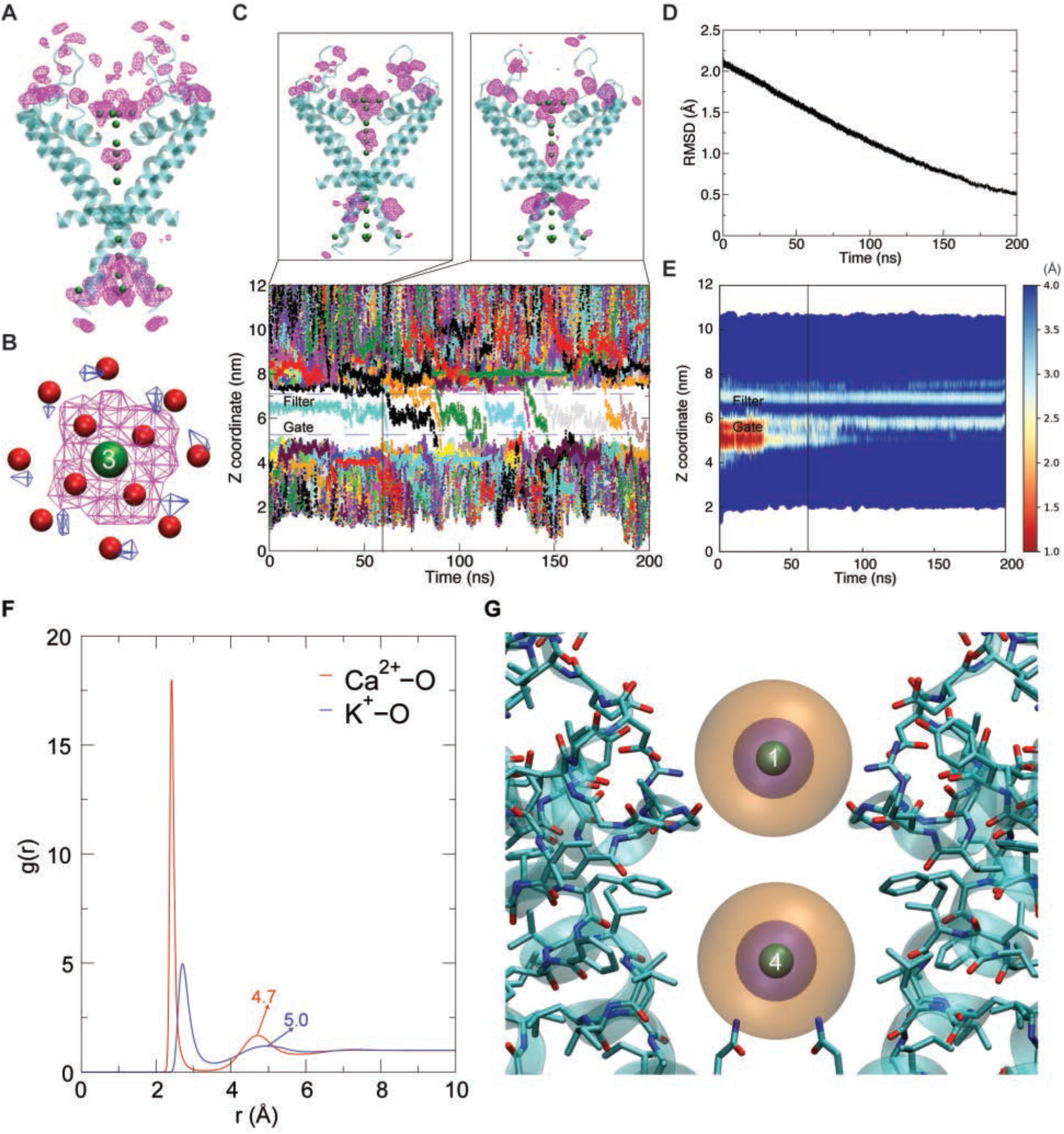
Classical and targeted molecular dynamics simulation of the Ca^2+^ permeation and interactions with RyR1. (**A**) The most probable binding sites of the Ca^2+^ ions are shown as magenta meshes in the MD simulation of state **1** without applying transmembrane voltage. (**B**) The most probable binding sites for eight stabilized water molecules at the vertical position of Site 3 observed in the MD simulations are shown as blue meshes and are superimposed with the water molecules in the atomic model determined from the cryo-EM density map of state **1**. The positional difference of the eight water molecules between the MD simulation and cryo-EM density peaks is in the range of 0.6-1.2 Å, much below the measured 2.6-Å resolution of the cryo-EM map. This indicates a remarkable agreement between the MD simulation and cryo-EM reconstruction on the stabilized locations of these water molecules in the central cavity. (**C**) The Ca^2+^ distribution and permeation through the gradually opened RyR channel in the targeted MD simulation. The Ca^2+^ ions started to permeate at around 60 ns in our targeted MD simulations in the lower panel, where the z coordinates of the Ca^2+^ were shown with different colored lines. The most probable binding sites of the calcium ions before and after permeation are shown as magenta meshes in the upper panels. Only two chains on the opposite sides from the cryo-EM structure were shown for clarity, and the Ca^2+^ ions from the cryo-EM density map were shown as green spheres. (**D**) The RMSD of the protein backbone during the 200-ns targeted MD simulation, the open-state structure being the reference (RMSD = 0). (**E**) The evolution of the pore radius during the 200-ns targeted MD simulation, showing the pore was gradually opened. (**F**) The radial distribution functions (RDFs) of water oxygen atoms around cations from our MD simulations. The first (left) and second (right) peaks correspond to the inner and outer hydration shells of water molecules, respectively. The peak position of the second hydration shell are labelled. (**G**) The size of the first (purple sphere) and second (orange sphere) hydration shells of the Ca^2+^ ions at Sites 1 and 4 (green sphere). Only two opposite protein chains are shown for clarity.

**Fig. S8.**
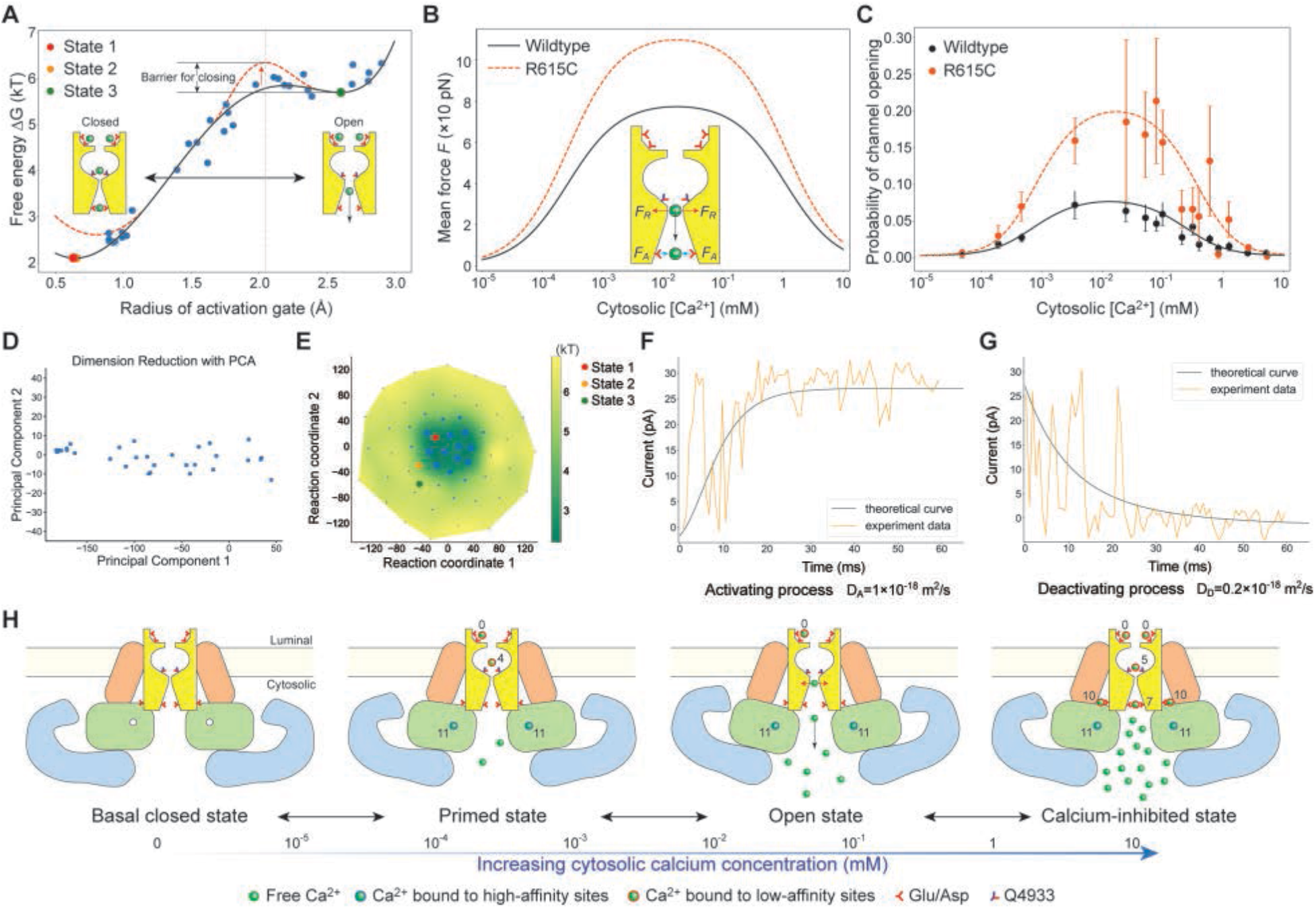
Energy landscape and stochastic dynamics simulations of the nonequilibrium gating of RyR1. (**A**) Experimental free-energy profile plotted against the radius of activation gate (black curve), fitted from the free-energy data points (blue filled circles) of 33 cryo-EM density maps estimated using the Boltzmann distribution. The red dashed curve shows the putative free-energy profile of RyR1 modified with permeating Ca^2+^ flux based on single-channel experiments^41^. (**B**) The simulated mean force imposed on the channel plotted against the cytosolic Ca^2+^ concentration for wild-type RyR1 (black solid) and R615 mutant^41^ (orange dashed). *F*_R_, repulsion force. *F*_A_, attraction force. (**C**) Cytosolic Ca^2+^ response curve of wild-type RyR1 (black solid) and R615 mutant (orange dashed). The results from our SD simulations are plotted as solid curves in agreement with experimental data (dots with error bars) in the literature^41^. (**D**) Mapping of 33 density maps to two dimensions using linear dimensionality reduction by principal component analysis (PCA). (**E**) Mapping of 53 density maps to two dimensions using nonlinear dimensionality reduction by t-distributed stochastic neighbor embedding (t-SNE). (**F** and **G**) Kinetics of channel opening (**F**) and closing (**G**) from single-channel experiments (orange) and from our SD simulations (black). (**H**) Graphic summary of the mechanism of Ca^2+^-regulated activation and inhibition of the RyR1 activity and associated conformational changes.

**Table S1.** Cryo-EM data collection, structure determination and refinement statistics.

|  | State 1 | State 2 | State 3 |
| --- | --- | --- | --- |
| **Data collection and processing** |  |  |  |
| Magnification | 105,000 | 105,000 | 105,000 |
| Voltage (kV) | 300 | 300 | 300 |
| Electron exposure (e-/Å<sup>2</sup>) | 49 | 49 | 49 |
| Defocus range (μm) | -0.6 to -3.5 | -0.6 to -3.5 | -0.6 to -3.5 |
| Pixel size (Å) | 0.685 | 0.685 | 0.685 |
| Symmetry imposed | C4 | C4 | C4 |
| Initial particle images (no.) | 835,709 | 835,709 | 835,709 |
| Final particle images (no.) | 389,292 | 112,781 | 3,922 |
| Map resolution (Å) | 2.6 | 3.3 | 6.3 |
| FSC threshold | 0.143 | 0.143 | 0.143 |
| Map resolution range (Å) | 2.5-4.8 | 2.5-5.8 | 4.5-12.0 |
| **Refinement** |  |  |  |
| Initial model used | 5T15 |  | 5TAL |
| Model resolution (Å) | 3.4 | 3.8 | 7.8 |
| FSC threshold | 0.5 | 0.5 | 0.5 |
| Model resolution range (Å) | 2.5-4.8 | 2.5-5.8 | 4.5-12.0 |
| Map sharpening *B* factor (Å<sup>2</sup>) | -50 | -40 | -46 |
| **Model composition** |  |  |  |
| Non-hydrogen atoms | 117,618 | 117,571 | 117,500 |
| Protein residues | 16,680 | 16,680 | 16,672 |
| Ligands | 70 | 23 | 8 |
| *B* factors (Å<sup>2</sup>) |  |  |  |
| Protein | 23.2-216.42 | 30-206.57 |  |
| Ligand | 32.83-192.67 | 64.5-128.59 |  |
| **R.m.s. deviations** |  |  |  |
| Bond lengths (Å) | 0.008 | 0.007 | 0.008 |
| Bond angles (°) | 1.065 | 0.994 | 1.154 |
| **Validation** |  |  |  |
| MolProbity score | 2.6 | 2.8 | 2.9 |
| Clashscore | 5.13 | 5.53 | 6.13 |
| Poor rotamers (%) | 0.52 | 0.3 | 0.4 |
| **Ramachandran plot** |  |  |  |
| Favored (%) | 89.07 | 88.60 | 89.17 |
| Allowed (%) | 10.65 | 11.25 | 10.68 |
| Disallowed (%) | 0.29 | 0.15 | 0.15 |

**Table S2.** Model parameters used in stochastic dynamics simulations.

| Parameter | Value |
| --- | --- |
| Area tension of channel ($\alpha$) | $0.120 \text{ J m}^{-2}$ |
| Radius of the stable state ($R_0$) | $0.645 \text{ \AA}$ |
| Temperature ($T$) | $310 \text{ K}$ |
| Mean hydrophobic force of single $\text{Ca}^{2+}$ ion ($f_1$) | $1.995 \times 10^{-11} \text{ N}$ |
| Mean electrostatic interaction of single $\text{Ca}^{2+}$ ion ($f_2$) | $1.995 \times 10^{-11} \text{ N}$ |
| Local particle number in the channel ($N$) | $4$ |
| Association constant 1 ($\kappa_1$) | $4 \times 10^6 \text{ M}^{-1}$ |
| Association constant 2 ($\kappa_2$) | $1.45 \times 10^3 \text{ M}^{-1}$ |
| Gate radius threshold releasing calcium ($R^*$) | $2.74 \text{ \AA}$ |

**Movie S1**. 3D Motion illustration of overview on the 21 Ca^2+^-binding sites of the RyR1 structure in state **1**. The Ca^2+^ ions are shown as green spheres.

**Movie S2**. A movie directly recorded in the complete procedure of the 200-ns targeted MD simulations from the closed to open state shows single-file Ca^2+^ flux, its interactions with the pore and the permeation behavior. The Ca^2+^ ions are shown as green spheres.

**Movie S3**. 3D motion illustration of the Ca^2+^ hydration complex bound at Site 1 in the selectivity filter. The Ca^2+^ ions and oxygen atoms of water molecules are shown as green and red spheres, respectively. The first hydration shell is highlighted by blue dashed lines connecting the water molecules, whereas the green dashed lines highlight the potential hydrogen bonds between the first or second hydration shells and the carbonyl or carboxyl oxygen atoms in the selectivity filter.

**Movie S4**. 3D motion illustration of the Ca^2+^ hydration complex bound at Site 4 in the central cavity, corresponding to a plausible occupancy state observed in our MD simulations, in which Sites, 2, 3 and 5 are occupied with water molecules, and Site 1 by a Ca^2+^ ion. The Ca^2+^ ions and oxygen atoms of water molecules are shown as green and red spheres, respectively. The first hydration shell is highlighted by blue dashed lines connecting the water molecules, whereas the green dashed lines highlight the geometry of the second hydration shell.

**Movie S5**. 3D motion illustration of the Ca^2+^ hydration complex bound at Site 7. The Ca^2+^ ions and oxygen atoms of water molecules are shown as green and red spheres, respectively. The first hydration shell is highlighted by blue dashed lines connecting the water molecules, whereas the green dashed lines highlight the potential hydrogen bonds between the hydration shells and the side-chain oxygen atoms in the cytosolic vestibule.

